# Cascading effects of species loss on ecosystem service provisioning in a marine food web

**DOI:** 10.1101/2025.10.20.683080

**Authors:** Anna Eklöf, Johanna Sollén Mattsson, György Barabás

## Abstract

Ecosystem services are important for the well-being of human societies. At the same time, the ecosystems providing the services are under constant pressure from anthropogenic activities, causing loss of biodiversity. Since all species are connected to each other via interaction networks, the loss of one species can trigger additional, secondary extinctions. However, the consequences of these secondary extinctions for ecosystem services are not known. Here we address this gap by coupling threats to species and their interaction networks, and these networks to ecosystem service provisioning. We use a highly resolved food web dataset of the Barents Sea, and simulate secondary extinctions using Bayesian networks. We explore a range of assumptions on how the loss of service-providing species translates to the disintegration of the services they provide.

We find that activation of threats cause secondary extinctions of service-providing as well as supporting species which negatively affect ecosystem services, and that the magnitude of these effects are highly dependent on the sensitivity of consumers to resource loss as well as service robustness. We also show that the indirect effects from species interactions causes substantial decreases in service provisioning. Understanding the linkages between ecological threats, species interactions in ecological networks and ecosystem service provisioning will be an important tool for guiding management strategies for effective ecosystem service management.

## Introduction

Loss of biodiversity is one of the most severe threats the Earth is facing today (Cardinale *et al*., 2012) as numerous threats from human activities are causing rapid decline of species from a wide spectrum of taxonomic groups (Díaz *et al*., 2019). Threats are for example over-harvesting, pollution and eutrophication (Maxwell *et al*., 2016; Tilman *et al*., 2017; Holma *et al*., 2019). At the same time, human societies depend on goods and services originating from the species in the ecosystems, for example provisioning of food, carbon sequestration and coastal protection. Some services are provided by single species, so if that species goes extinct the service will naturally also disappear. A service can also be provided by groups of species, and in those cases the loss of a service is likely a function of number of species that are service providers (Dee *et al*., 2017a).

Species in ecosystems are connected to each other forming complex networks of interactions crucial for their survival. Therefore, the loss of a species can cause additional (secondary) extinctions (Dunne *et al*., 2002; Eklöf & Ebenman, 2006). Even if the primary extinction was not of a service-providing species, species that are service providers in other, sometimes distant, parts of the network can go secondarily extinct leading to loss of important services. Nevertheless, although the network approach has been highlighted as one of the important research paths (Jacob *et al*., 2020; Dee *et al*., 2017a) the consequences of these secondary extinctions for ecosystem services have rarely been analyzed (but see Keyes *et al*. (2021)).

There are many different approaches to modeling secondary extinctions in ecosystems. A flexible and computationally effective method is provided by Bayesian networks. This method has gained recent traction in ecology as a way of modeling secondary extinctions (McDonald-Madden *et al*., 2016; Staniczenko *et al*., 2017; Häussler *et al*., 2020), especially as its predictions closely match those obtained from more complex and cumbersome methods (Eklöf *et al*., 2013). Bayesian networks follow the fundamental principle that a predator is more likely to go extinct if its prey are also more likely to go extinct. Thus, the percolation of extinction risk in a food web is modeled bottom up, using the law of total probability to obtain the marginal extinction likelihood of each species in the network (Eklöf *et al*., 2013). By highlighting species and links being especially important to extinction risk propagation, the method also provides insight into how conservation efforts could be directed to mitigate any adverse effects (McDonald-Madden *et al*., 2016).

Threats to ecosystems are multifaceted and depend on the nature of the disturbance. Specific threats may affect species differently. Some, like selective fishing for food, directly target specific species. Others, such as climate change, are more universal – though they still need not impact all species equally. Threats can act both on local (e.g., oil spills) and regional/global scales (climate change) (HELCOM, 2009). Additionally, different threats can also have synergistic effects, and as climate change continues to accelerate, threats are increasingly likely to exacerbate each other’s negative impacts on ecosystems (McComb & Cushman, 2020).

The outlined sequence of events (threat – species network – ecosystem services), implies that the effect of a disturbance on service provisioning can be both complex and non-intuitive. Therefore, in order to take well-founded conservation actions, we need a deeper understanding of how different threats and ecosystem services relate to different species and, in particular, the structure of their interactions in several steps (Jacob *et al*., 2020). This knowledge will allow for a more complete accounting of the total services provided and how sensitive they are to different disturbances.

Here, we examine how the activation of ecological threats propagates through species interaction networks to affect ecosystem service provisioning using a data set of the Barents Sea food web.(Kortsch *et al*., 2018). Like all ecosystems, the Barents Sea provides multiple benefits for the human societies. Example of services are food provisioning, recreation and coastal protection (Holma *et al*., 2019). Simultaneously, the Barents Sea is also exposed to numerous, severe environmental threats due to the intensity of the human activities in the area, as well as global threats. We evaluate how various threats of different magnitude (severity for different species) propagate through the ecological network of interacting species and further to ecosystem services in the Barents Sea. We use Bayesian networks to model how primary extinctions cause secondary extinctions. We then analyze how ecosystem service provisioning is affected by biodiversity loss depending on the identity and nature of the threat. This include evaluating difference in response dependent on i) species targeted, ii) ecosystem service targeted, and iii) spatial location of local food webs. We find that secondary extinction causes substantial declines in ecosystem service provisioning, in particular for services delivered by species at high trophic levels.

## Methods

### Data

We used a food web dataset describing the Barents Sea (Planque *et al*., 2014; Kortsch *et al*., 2018). The Barents Sea is a shelf sea with a heterogeneous environment, bordering the Atlantic Ocean with the dissipating Gulf Steam in the west, and the Arctic Ocean to the north-east (Fossheim *et al*., 2015). The food web data consist of a regional metaweb and 25 subregional local food webs. The subregions used in the Barents Sea dataset are delimited by polygons defined by a group of experts in Hansen *et al*. (2016), aiming to make the subregions as homogeneous as possible with regards to hydrography and bathymetry.

The metaweb includes 233 species and 2220 feeding interactions, of which species occurrences are based on catch data, and interactions are based on the ecological literature (Planque *et al*., 2014; Kortsch *et al*., 2018). Species range from primary producers to avian and mammalian top predators. The subregions include separate local species occurrences with 115–178 species. Interactions in each subregion were inferred from the metaweb any pair of taxa with an interaction in the metaweb was assumed to interact at any site where the two taxa co-occurred (Planque *et al*., 2014; Kortsch *et al*., 2018).

### Identifying threats and ecosystem services for analyses

We reviewed a wide range of scientific and gray literature to identify the major threats affecting organisms in the Barents Sea. To ensure consistency and comparability, we only retained threats that were reported across multiple species or taxonomic groups, and for which sufficient information was available to support subsequent threat scoring (see section Quantifying the impact of threats on individual taxa, for details SI). Threats mentioned exclusively for one or two species, or lacking adequate documentation, were excluded from further analysis.

The assessment was informed by several key sources, including *OSPAR’s List of Threatened and/or Declining Species & Habitats* (OSPAR, 2024) *and pages therein, Artsdatabanken’s 2021 Red List Assessments (Artsdatabanken, 2021), the IUCN Red List assessments (IUCN, 2023a)*,

*Barents Sea Ecoregion – Ecosystem overview* (ICES, 2021), *and the Joint Norwegian - Russian environmental status 2013. Report on the Barents Sea Ecosystem*. (McBride *et al*., 2016). For a full reference list, see SI.

We identified eight threats our analyses: 1) habitat destruction, 2) ice loss and increased water temperature, 3) pollution, including marine litter, 4) ocean acidification, 5) oil pollution, 6) pathogens, 7) seabed abrasion and 8) selective extraction of species. In turn, we analyzed the following ecosystem services: 1) carbon sequestration, 2) habitat provisioning, 3) food provisioning and 4) recreation and tourism (Table **??**). For a detailed description of the threats and ecosystem services, see SI.

### Mapping species to threats and ecosystem services

We conducted a literature search to compile information on how species in the Barents Sea are affected by threats and contribute to ecosystem services. The search was not intended as a systematic review in the strict methodological sense, but as a structured effort to gather a representative body of evidence. Google Scholar served as the primary database, with the acknowledged limitations of incomplete coverage, variable indexing, and limited reproducibility.

Specifically, we then searched for the unique species–threat (or species–ecosystem service) combination, using both species common and scientific names, as well as synonyms for each threat/service. For species that could still not be linked to threat/service, we looked at taxonomically similar species; if close relatives were service providers, the focal species was also considered to provide that service. Some species could be tied to habitats that were providing a service. In that case, the species–threat/service links were identified based on the habitat the species primarily belong to.

Both peer-reviewed journal articles and gray literature were included. Grey literature was defined as reports and documents issued by governmental authorities, research institutes, intergovernmental organizations, scientific commissions, and professional societies. Only sources explicitly addressing threats, species, or ecosystem services in Arctic or Barents Sea ecosystems, were retained for analysis.

If a threat was identified to affected a species, that species’ background extinction risk (see below) was increased. One can view this as adding a link from the threats to the respective species the threat is affecting. In a similar manner, when a species was identified as a service provider, we added a link from that species to the service. Thereby we formed a multilayer network (Kivelä *et al*., 2014; Pilosof *et al*., 2017) with three layers; threats, species and ecosystem services.

### Analyzing extinction probabilities using Bayesian Networks

In order to analyze if threats towards specific species cause increased probability of secondary extinctions of other species in the food web, we follow the Bayesian network approach outlined in Eklöf *et al*. (2013). Modeling species interactions as Bayesian networks allow the species extinction probabilities to increase gradually with resource loss following some pre-defined function. Additionally, the species extinction probabilities can be nonzero even when species have full access to their resources – all species have a baseline probability of extinction. This baseline probability quantifies the probability of species going extinct for causes other than those represented by the interaction network. Here we use this baseline probability of extinction to model how each species is affected by threats.

We model the probability *P* (*¬C*|*f*) of a species *C* going extinct as a function of the fraction

*f* of its resources being absent:

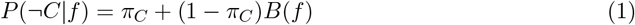

where *π*_*C*_ is species *C*’s baseline extinction probability, and *B*(*f*) is a monotonically increasing function of *f* such that *B*(0) = 0 and *B*(1) = 1. For any basal species *R*, we assume that all of its resources are always present, so *f* = 0 and *P* (*¬R*) = *π*_*R*_ is simply its baseline extinction probability. For a non-basal (consumer) species *C*, one obtains *P* (*¬C*) by using *P* (*R*) = 1 *− P* (*¬R*) and the law of total probability. For instance, if *C* has two prey items *R*_1_ and *R*_2_, we write

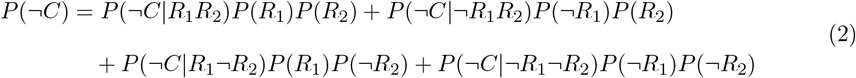

where *P* (*¬C*|*R*_1_*¬R*_2_) is the probability of *C* being extinct given that *R*_1_ is extant and *R*_2_ is extinct, and so on. Knowing *P* (*¬R*_1_) and *P* (*¬R*_2_) (either because they are basal species, or the same formula has been already used to derive their values), we can then calculate *P* (*¬C*) from Eq. 2. Thus, determining the extinction probabilities of all species in a food web is a bottom-up calculation process: we start with basal species, then move on to species only consuming those basal species, and so on.

We can vary how sensitive consumers are to losing resources in the Bayesian network by varying the functional form of *B*(*f*). Here we implement three different forms of this response function:

- *B*(*f*) = *f* (linear consumer). The probability that the consumer goes extinct is directly proportional to the fraction of resource species it has lost.
- *B*(*f*) = *f* ^2^ (resilient consumer). The consumer’s extinction probability only starts appreciably increasing after some fraction of the resource species have already been lost.
- *B*(*f*) = 1 *−* (1 *− f*)^2^ (sensitive consumer). The probability that the consumer goes extinct attains high values already after the loss of a small fraction of its resources.

### Quantifying the impact of threats on individual taxa

As different threats are can give more or less severe effects, which also varies depending on the species exposed. We therefore used a standardized method for determining the *magnitude* of the different threats analyzed.

We base our method on the IUCN threat impact scoring scheme (IUCN, 2023b), with some modifications. In our approach, each threat is assessed using two factors — scope and severity — each assigned a value of 1, 2, or 3. These two scores are then added together.

Both scope and severity can also take the value “Unknown”, indicating that the threat is expected to affect the species, but the extent of that effect is uncertain. This differs from cases where a species is not directly affected by the threat, for which the score is 0.

To account for the “Unknown” category, we normalize our scoring so that a species with both scope and severity set to Unknown receives a score of 1. As a result, scores range from 1–7 for affected species and 0 for unaffected species.

These scores are then translated into extinction probabilities ranging from 0.1 – 0.7. In addition, we assume all species face a background extinction risk (unrelated to the specific threat) of 0.1. Therefore, the total extinction risk for a species ranges from 0.1 (for unaffected species) to 0.8 (for species affected by a threat with maximum scope and severity, i.e., 3 each) For details, see SI.

### Calculating the loss of ecosystem services

Ecosystem services are provided by single or, more often, several species. This means that the loss of ecosystem service provisioning depends on which and how many of the service-providing species are primarily and secondarily lost in the food web. We do the mapping from species losses to ecosystem service losses using a summary function *σ*(*p*_*i*_) that takes the vector of species’ persistence probabilities *p*_*i*_ as input and gives a single value between 0 and 1 as its output, interpreted as the fraction of the original ecosystem service provision that is retained after a threat is realized. To explore how results depend on the functional form of *σ*, we use three different choices:

1. 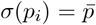(proportional ESS), where 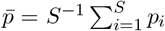is the mean persistence probability of all *S* species in the system. The fraction of services lost thus depends directly and linearly on the extinction risks of the service providing species.
2. 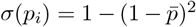(robust ESS). The fraction of the service lost increases slowly at first, and starts appreciably degrading only after some amount of the service-providing species have been lost.
3. 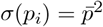(vulnerable ESS). Here the fraction of the services being lost increases sharply already when a small fraction of the service providing species have been lost.

We additionally analyzed more pronounced versions of the robust and vulnerable summary functions as well (SI).

### Effects of species interactions

The mapping of threats to ecosystem service loss can be more or less affected by the species interactions in the network. In cases where the species targeted by the threat are exactly the same species providing the service, the effects of network structure will be unimportant. On the other hand, when species targeted by a specific threat differ from the ones providing the service in focus, species interactions are potentially important for the effects on ecosystem service provisioning. Therefore, we analyzed the overlap between threatened species and service-providing ones. For each subregion, and for all combinations of threats and ecosystem services, we identified the number of species that are i) only affected by the threat, ii) only provide the ecosystem service, iii) are both targeted by the threat and provide the ecosystem service, and iv) are neither targeted by the threat nor are ecosystem service providers.

In addition to the number of species affected by a threat or providing ecosystem services, the trophic level of these species can strongly influence how the activation of different threats propagates through the ecosystem, particularly given our bottom-up approach in which consumer persistence depends on the availability of their resources. Accordingly, we quantified the trophic positions of both threatened and service-providing species.

To evaluate the importance of the network structure for transferring the threat to ecosystem service provisioning, we also ran our analyses without taking the network structure into account. This was done by simply neglecting all species interactions.

## Results

The degree of ecosystem service retention depends both on which threat is activated and the specific service in focus. Habitat provisioning and Carbon sequestration are the services exhibiting the smallest relative losses, whereas Food provisioning and Recreation and tourism experience the greatest reductions, across all threats (Fig. 1).

**Figure 1:**
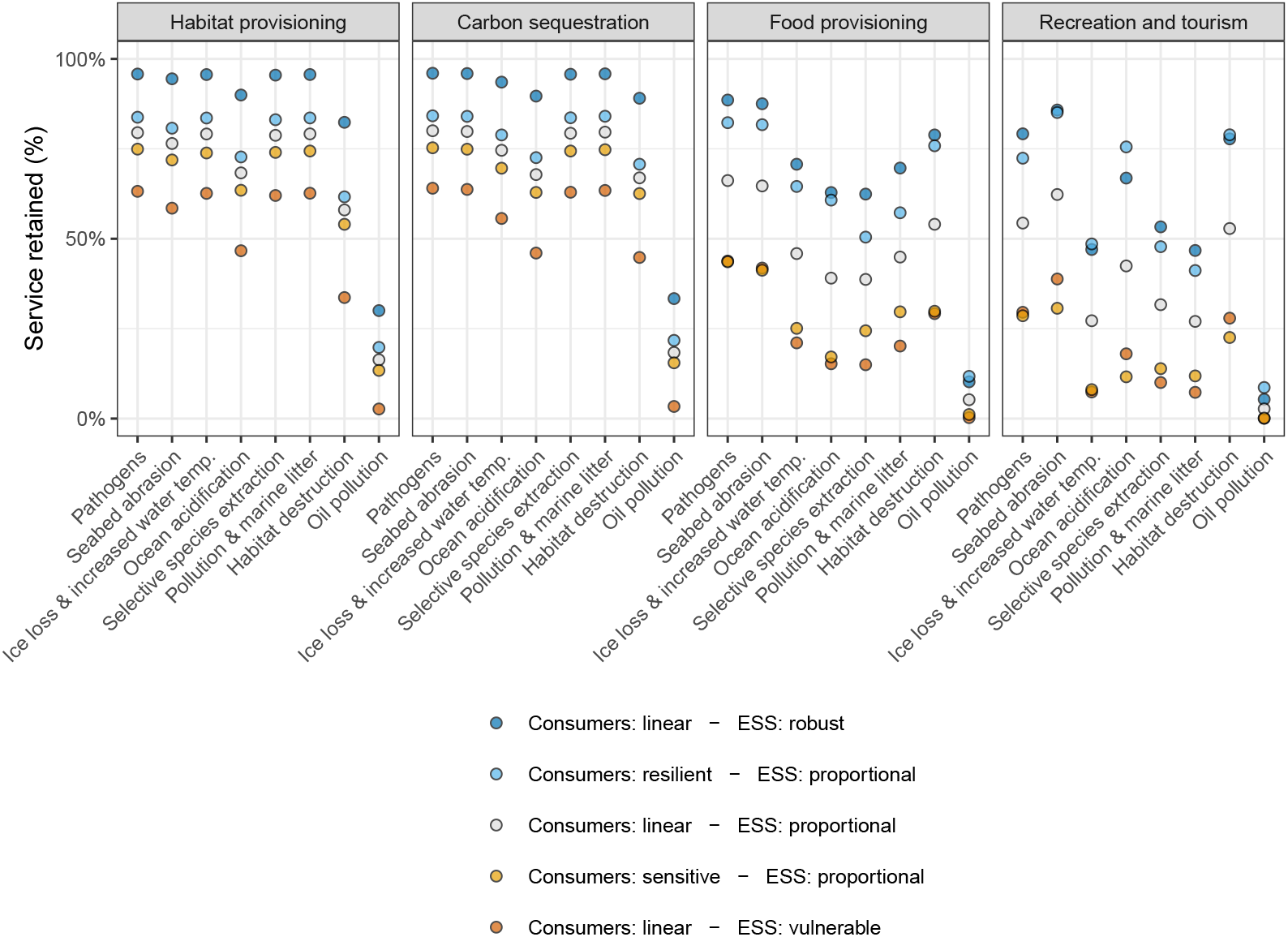
Fraction of ecosystem services retained in subregion 46 of the Barents Sea (y-axis) for different ecosystem services (panels) when different threats (x-axis) are realized. Results are shown for five different combinations of consumer response and ecosystem service summary functions (colors; see Methods). The general trends are similar across all other subregions as well (SI).

Pathogens and Sea abrasion are the two threats causing the smallest reduction in ecosystem services across all services, with more than 75% retention for Habitat provisioning and Carbon sequestration and 60-70% for Food provisioning and Recreation and tourism in the scenario with linear consumer responses and proportional reduction in ecosystem services (Fig. 1). Oil pollution, on the other hand, gives the largest overall losses. Carbon sequestration and Habitat provisioning preserve only about 20% of their original values while Food provisioning and Recreation and tourism, fall below 10% even in the most favorable scenario, underscoring that oil pollution drives a uniformly sharp decline in ecosystem service provisioning. For the remaining threats, the pattern is more service-specific. For instance, activation of the threat of Ice-cover loss results in a retention level of 27% for Recreation and tourism but over 75% for Habitat provisioning (again for linear consumer responses and proportionally-reducing ecosystem services). Carbon sequestration is only slightly more affected than habitat provisioning, while Food provisioning is retained at about 48%.

The effects of the different threats on ecosystem service retention depend strongly on both the consumer response to prey loss and the summary function converting species loss to ecosystem service reduction (Fig. 1). For simplicity, in our analyses we keep one of the functions linear/proportional, and vary either the consumer response or the ecosystem service summary function.

For Carbon sequestration and Habitat provisioning, the ordering of the functions is coherent; across all threats, service retention consistently increases along the gradient from consumer linear - service vulnerable, consumer sensitive - service proportional, consumer linear - service proportional, consumer resilient – service proportional, to consumer linear – service robust (Fig. 1). By contrast, the patterns for Food provisioning and Recreation and tourism are more variable; the two functions giving the highest and lowest service retention, respectively, are in majority of threats the same but their internal ordering varies depending on which threat is activated.

### Differences between subregions

In addition to being service-specific, the fraction of ecosystem services that remain after a threat is activated differs across subregions. For the Ice loss threat, for example, the services Carbon sequestration, Food provisioning and Recreation and tourism exhibit pronounced differences in retention between subregions (e.g., Recreation and tourism retention ranges from 56% to 27%, Fig. 2). For the service Habitat provisioning, the difference in Ice loss retention subregions between is more limited. For other threats such as Seabed abrasion, the difference in response between subregions is less pronounced across all services.

**Figure 2:**
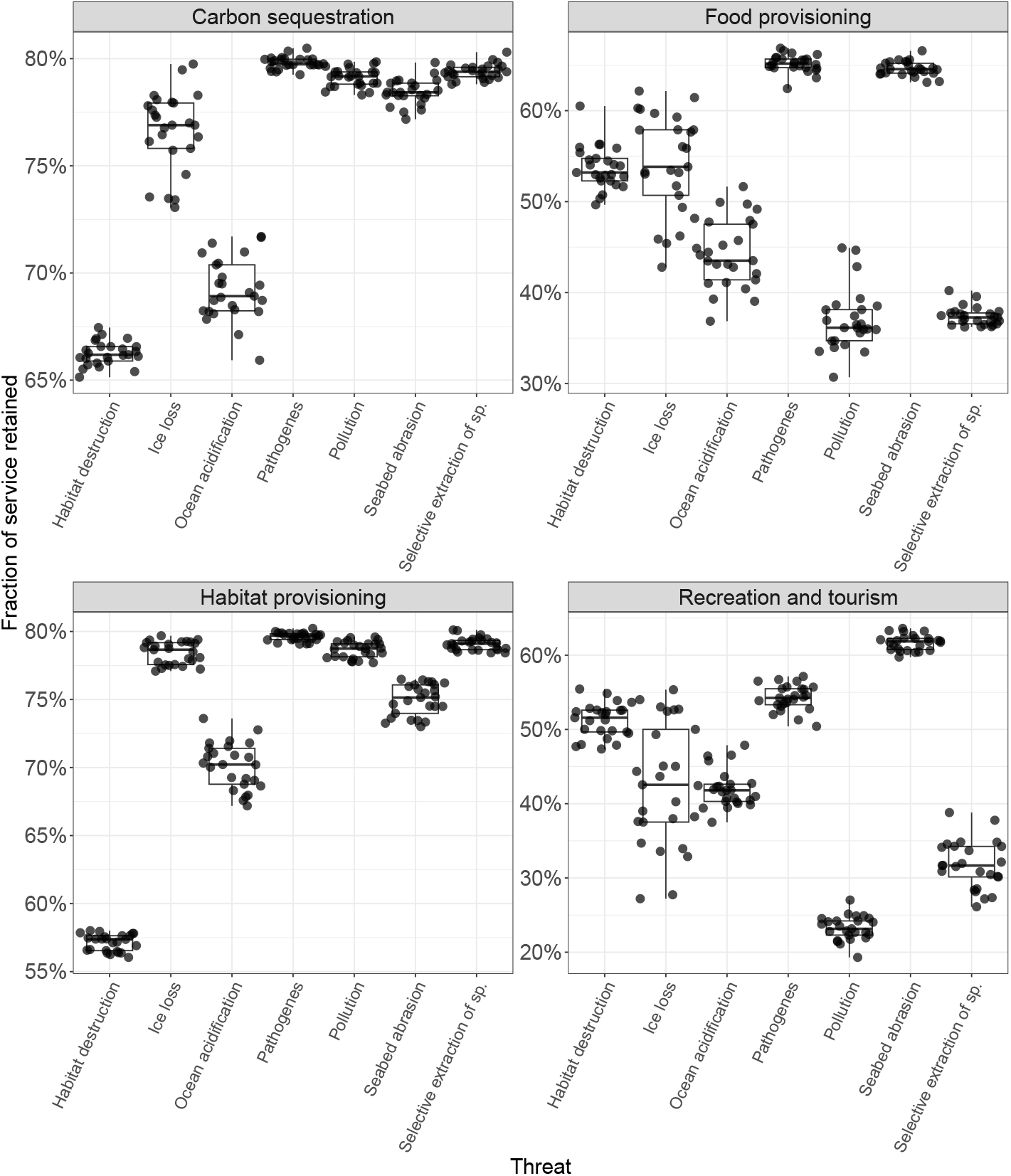
The fraction of different ecosystem services retained in the different subregions (points) after various threats (panels) are realized. The box plots summarize results from all subregions. (Boxes go from the first to the third quartile, the horizontal line shows the median, and the whiskers the minimum and maximum values.) The x-axis shows the ecosystem service analyzed. They are from simulations with linear consumer response and proportional ecosystem service summary functions. To improve visibility, Oil pollution is excluded since its much lower values would compress the scale and obscure variation among subregions for the other threats (the y-axes differ between panels).

The threats differ both in the number of species being threatened and their magnitude (Fig. 3). Oil pollution stand out as the threat affecting many species and also having a high magnitude. On the opposite end of the scale is Pathogens, which are affecting few species to a relatively low magnitude. There are also differences in both magnitude and number of species affected between subregions; For example, for Ice loss, the average magnitudes varies between 0.6 and 0.74. For Pollution the number of species affected varies between 35 and 75 species.

**Figure 3:**
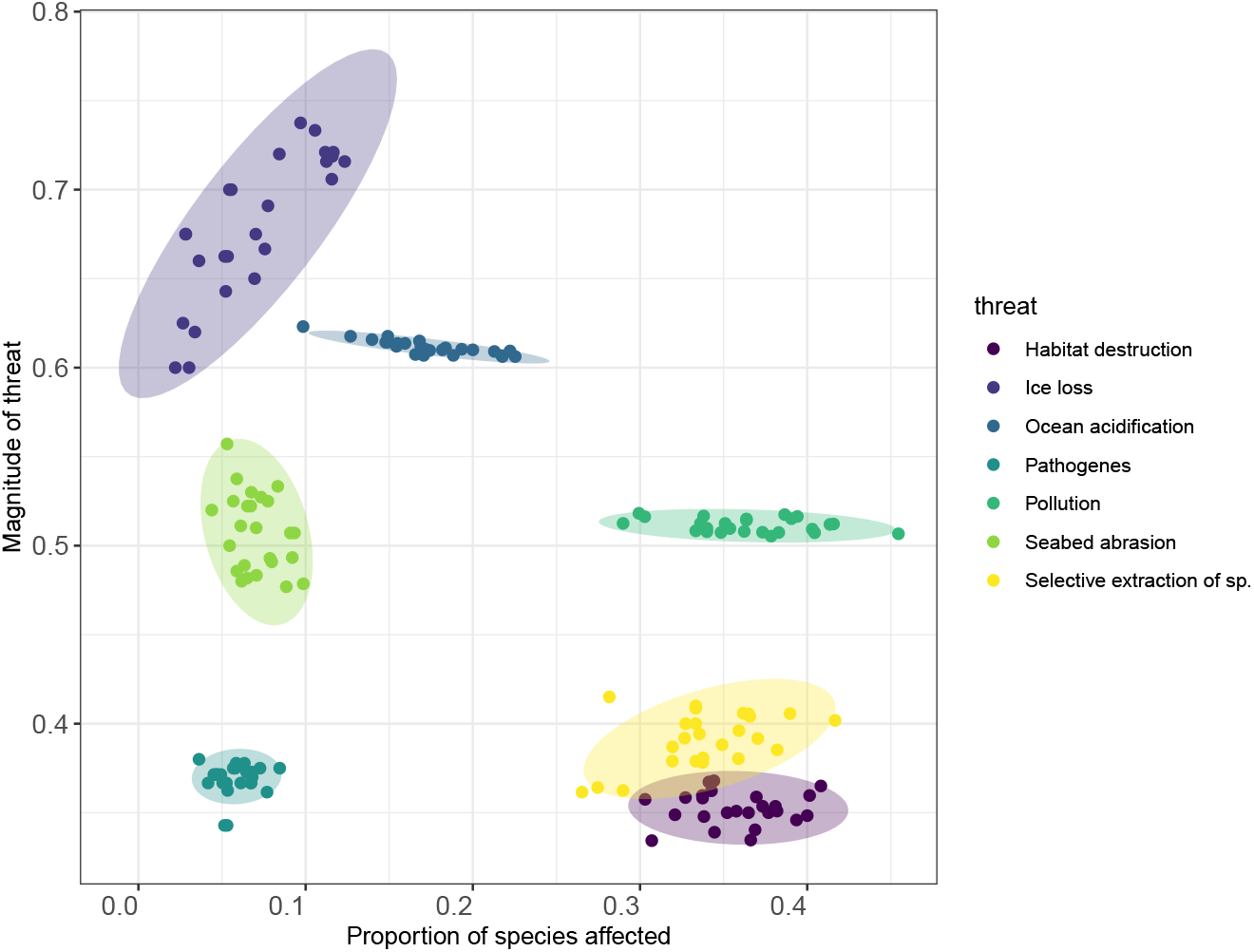
Proprtion of species affected (x-axis) by respective threats (colors), plotted against the average magnitude for all affected species (y-axis) in different subregions (points). Ellipses around points are for visual aid only, highlighting where the data fall.

### Direct versus indirect effects

If the species affected by a threat are the same as those providing the services, the effects from species interactions would be unimportant. However, for most combinations of threats and ecosystem services, there are discrepancies between species roles (whether a species is a service provider, affected by a threat, both, or neither). Fig. 4 shows the division of roles for the species in one subregion. (The division of roles is similar across subregions.)

**Figure 4:**
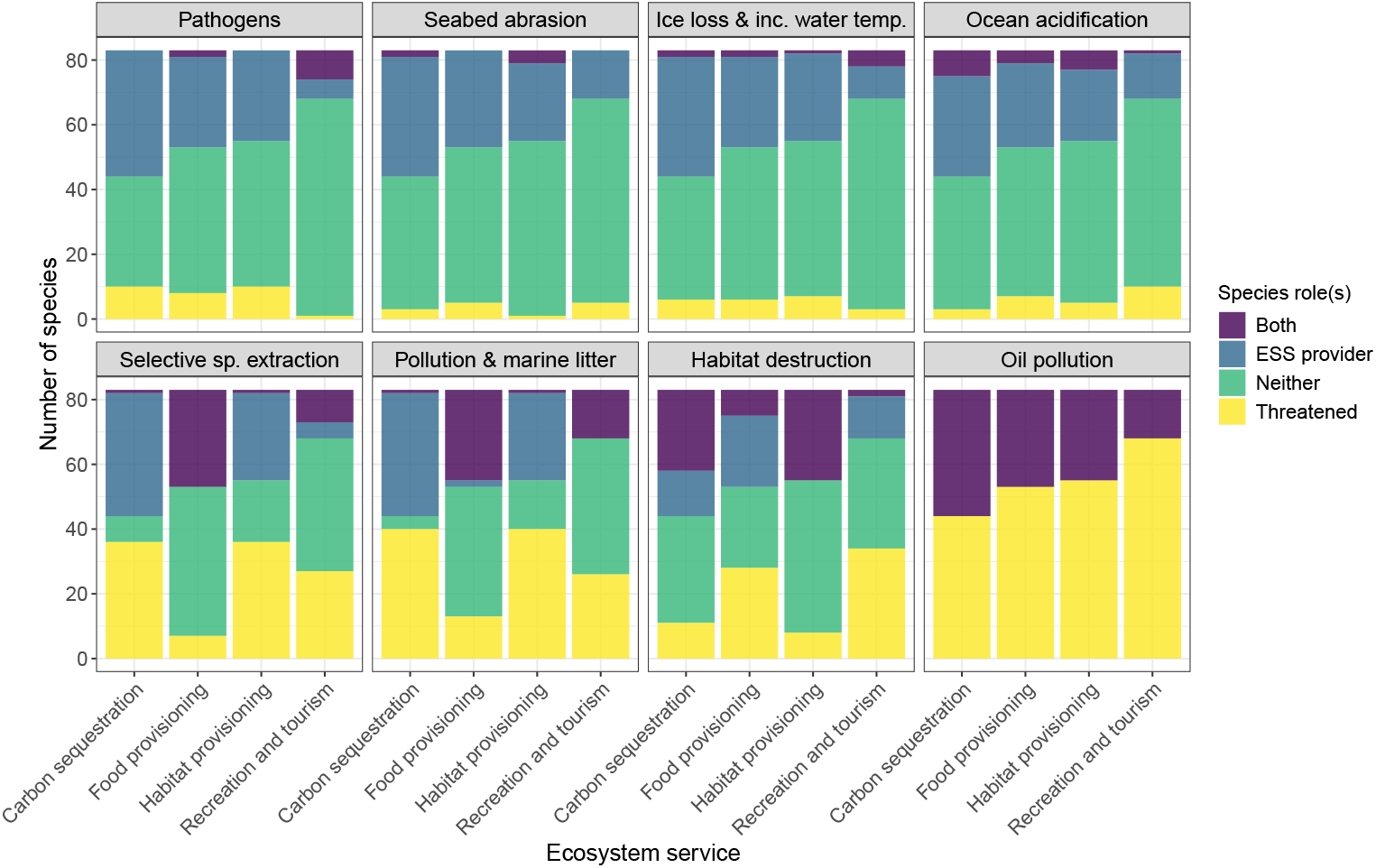
The number of species for respective combination of threat (panels) and ecosystem service (x-axis) that are either an ecosystem service provider (blue), directly affected by the threat (yellow), both a service provider and directly affected by the threat (purple), or neither a service provider or directly affected by the threat (green). Results are shown for one subregion (5) of the Barents Sea dataset.

For example, in the case of ice loss, only a few species are simultaneously threatened and serve as providers of ecosystem services, irrespective of the services considered. But for Selective extraction of species, a larger proportion of the species directly affected by the threat are also providing the service of Food provisioning (Fig. 4). However, across most threat–service combinations, species that are both directly affected and are also service providers are uncommon, except for Oil pollution.

A species’ position in the food web will affect its vulnerability to indirect effects. It is therefore important to analyze the distribution of trophic levels for threatened and service-providing species, respectively. The distribution of trophic levels for service-providing species varies across different services (Fig. S2). Habitat provisioning and Carbon sequestration are primarily provided by species at low trophic levels (Fig.S2a, b), whereas Food provisioning and in particular Recreation and tourism are provided mostly by species at higher trophic levels (Fig. S2c, d).

The distribution of trophic levels of species targeted by the different threats also varies, although the differences are less pronounced compared to the services. Some threats are nevertheless more specifically targeting species at certain trophic levels; e.g. for Selective extraction of species a majority of the targeted species are at high trophic levels, and for Habitat destruction most of the affected species are at low trophic levels (Fig. S3).

### Effects of species interactions

For every threat, incorporating the species interaction network markedly amplifies the negative impact activation of threats have on ecosystem-service provisioning (Fig.5). The magnitude of this amplification depends on the threat activated, the ecosystem service in focus and the different combinations of consumer response and ecosystem service summary. Across all threats except Oil pollution, sensitive consumers give the largest increase in service loss when network effects are taken into account (Fig.5), whereas resilient consumers give the smallest increase (Fig.5, yellow bars). The network effects for the other combination of functions for combinations of consumer response and ecosystem service summary functions are in between these two extremes.

**Figure 5:**
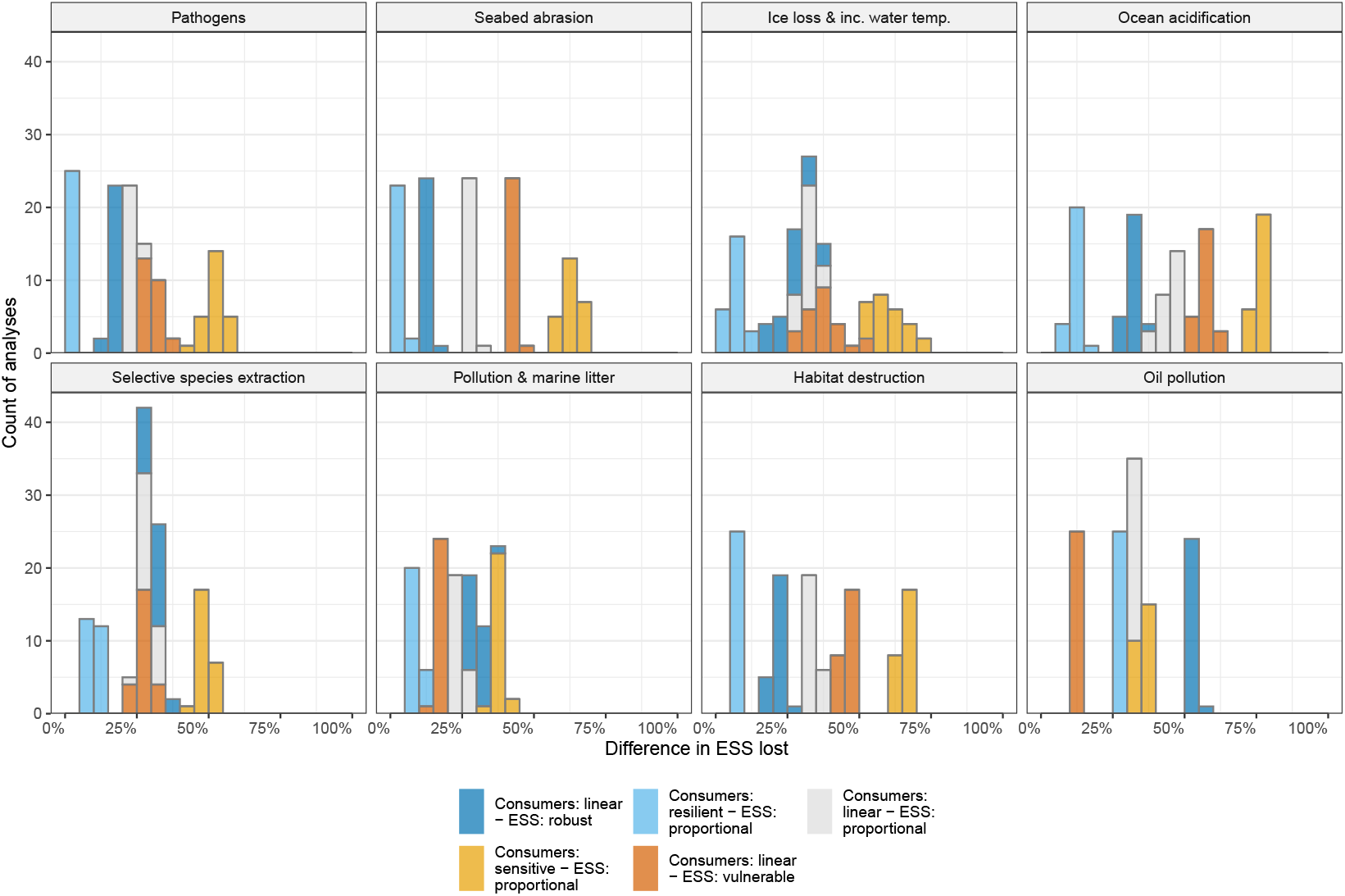
The number of analyses (y-axis) where indirect network effects caused increased loss of the ecosystem service Recreation and tourism after realization of different threats (panels). The x-axis shows the difference between ecosystem service retained when the species interactions are not taken into account versus when the interactions are taken into account, in percentages points. The histograms summarize calculations over the different subregions. Colors represent combinations of different consumer response and ecosystem service summary functions.

For Recreation and tourism with sensitive consumers, the network-mediated effect ranges from a 40% increase under pollution to a 75% increase under ocean-acidification. A similar range is observed for Food Production (see Fig S4). By contrast, Carbon sequestration and Habitat provisioning exhibit less variation among threats, with the maximum network-induced increase in service loss not exceeding approximately 25% (see Fig. S5 and S6).

## Discussion

We found that the retention of ecosystem services in the Barents Sea is greatly affected by the realization of different threats, with the magnitude of impact depending on both the identity of the threat and the service considered. In some cases, there is a direct mapping between threats and services, for example when the same species both experiences the threat and provides the service. However, indirect effects are often substantial.

These findings highlight two important points. First, there is often a disconnection between the species directly affected by threats and the species that act as service providers. Second, the vulnerability of ecosystem services may be greater than anticipated because many species indirectly support their provision. Recognizing these dynamics is crucial for effective ecosystem management.

Ecosystem services are often provided by groups of species, but different species relative contribution to the service is difficult to assess. It is likely that different ecosystem functions, and in the extension ecosystem services, have different sensitivities to loss of species; i.e. a service can be largely disrupted after only a few providing species been lost, or could be more resistant and stay retained following substantial losses Lyons *et al*. (2005); Smith & Knapp (2003). Here, we addressed these alternative possibilities by applying ecologically plausible functions to quantify how losses of service-providing species translate into service losses. As expected, all services were highly sensitive to the choice of functional response.

In this analysis, we used a generalized approach, assuming that all services shared the same functional response. However, in reality, some services are likely to exhibit more specific and characteristic responses. For example, services directly dependent on the extraction of specific species – such as Food provisioning – may be especially sensitive to the loss of key provider species (e.g., cod), whose species-specific contributions can in some cases be quantified (Dee *et al*., 2017b). For most services, however, such information remains limited. For habitat provisioning, for instance, relationships are often complex, depending on both the abundance and identity of contributing species (Winfree *et al*., 2015; Davies *et al*., 2011). The generic approach adopted here can therefore serve as a baseline that can be refined in future work by weighting species according to their relative contributions to a given service, for instance based on abundance (Xiao *et al*., 2018).

The same reasoning holds for consumer’s response to loss of resources. Some consumers may experience drastic declines already when a small fraction of its resources are lost, while other consumers might be able to withstand extensive losses. In our results we see, in some cases drastic, effects depending on how consumer’s response is modeled. For the services Habitat provisioning and Carbon sequestration changes in consumers’ sensitivity is causing less limited effects on service retention. However, for Food provisioning and Recreation and tourism there are substantial effects. For the threat Ocean acidification, for example, the largest differences in service retention is between consumers being resilient or sensitive to loss of resources. This is also reflected in the magnitude of network effects, as when consumers are sensitive the retention of Recreation and tourism decrease with more than 75% when species interactions are taken into account compared to when disregarded.

In general, the ecosystem services that are primarily provided by species at low trophic levels (Carbon sequestration and Habitat provisioning) are more robust to species losses than others meaning that they are retained to a higher extent, compared to services provided by species at high trophic levels. This is reasonable since when a service is provided for by lower trophic levels, the threat needs to also directly target those species in order to affect ecosystem service provisioning, given the bottom-up approach we use here.

For services primarily provided by species at high trophic levels, the relationship is reversed as a threat acting on low trophic level species percolates up in the food web with the possibility to have secondary effects on all trophic levels above. The increased extinction risk for species at higher trophic levels is in agreement with several previous studies (Eklöf & Ebenman, 2006; Curtsdotter *et al*., 2011; Ryser *et al*., 2019), and it is reasonable these losses cause increased risk of loosing ecosystem services.

However, these results reveal one of the limitations to the Bayesian Network approach; since it works in a strict bottom-up direction top-down effect are not taken into account. This means that we here likely underestimate the negative effects of biodiversity losses on ecosystem services.

The reasoning behind using this approach is that a fully dynamical simulation (i.e., coupled differential equations) of a highly complex network, such as the Barents Sea food web, would be impossible to parameterize to acceptable accuracy, and computationally demanding to run. The Bayesian network approach is also a substantial improvement compared to pure topological approaches (Eklöf *et al*., 2013; Häussler *et al*., 2020). In addition, it has earlier been shown that the Bayesian Network approach captured a majority of the secondary extinctions a fully dynamical model produce Eklöf *et al*. (2013), making this a reasonable option. That said, development of models for including top-down effects are still essential for more realistic evaluations of all indirect effects in ecological networks.

Although the network approach in ecosystem service analyses are still rare, there are some exceptions. Keyes *et al*. (2021) investigated how the ordering of species removals affected robustness of ecosystem services in three estuarine salt marches food webs. They showed that robustness differed dependent on which sequence that were executed, and their results revealed the importance of the species supporting the service providers. For example, removal of ecosystem service providers caused collapse of the ecosystem services, but not the full food web, whereas removal of supporting species additionally caused the food web to collapse and therefore had more far-reaching consequences. This is in line with our findings and underscores the importance of species interactions and indirect effects. Two important differences between our and Keyes *et al*. (2021) analyzes are that they used strictly topological simulations, whereas we take a more flexible approach. Also, while Keyes *et al*. (2021) perform specific removal sequences, we in our analyses take all removal sequences into account simultaneously and average over them. The two approaches have their advantages and disadvantages. For example, the Bayesian network approach include also sequences that are not predefined which mean the analyses do not miss out on potentially important extinction scenarios. On the other hand, the sequence approach can give detailed answers for a specific sequence, which might be of particular interest.

The dataset we use includes 25 different locations spanning the whole Barents Sea with its different environmental conditions. For some combinations of threats and services the fraction of service retained varied quite drastically between subregions (e.g., Ice loss – Recreation and Tourism and Increase water temperature – Food provisioning) the discrepancy between subregions were for some combinations limited, e.g., Seabed abrasion for all services. However, due to that the subregions are assembled from a metaweb (Kortsch *et al*., 2019) meaning that while species presence/absence in the different subregions was inferred based on several data sources (see Methods), the interactions were always taken from the metaweb. This means, that if two species have been determined to interact in any location they will do so in all local food webs where they both are present. In reality, this may not be the case as local conditions, such as presence of other species or environmental factors, can modify biotic interactions (Poisot *et al*., 2015). Therefore, it is likely that the differences in retained services between local food webs in fact are larger compared to the results presented here. This is a common problem in datasets of ecological networks Poisot *et al*. (2013). However, both empirical (Beng & Corlett, 2020) and theoretical methods (Cirtwill *et al*., 2019) are being developed to address this issue.

Here we have simulated each threat being activated in isolation. However, the various threats towards the Barents Sea ecosystem, or any ecosystem, does not need to be independent of one another. For example, Selective extraction of species can trigger also Seabed abrasion, depending on how the extraction is executed (e.g. bottom trawling). Various threats can potentially form a complex network in which a given threat may activate others. An interesting extension of our approach would be to construct a network of threats which by the couplings to the trophic network would form a multi-layer network (Pilosof *et al*., 2017) and the same analyses could be done.

Combining ecosystem service research with network research renders a pathway to understand the underlying role of species interactions for the direct as well as indirect effects that anthropogenic threats poses to ecosystem services. Our mathematical framework offers a flexible solution that can easily be extended as new data becomes available.

Our results pinpoint the importance of taking the network perspective when analyzing ecosystem service losses. To disregard species interactions can severely underestimate the negative effects from threats. Conservation actions targeting specific species without taking the interaction network into account can give unintentional outcomes. Our approach to combine threats, species interaction networks and ecosystem services in a multi-level network provides an important step for understanding how indirect effects of biodiversity losses causes risks for sustained ecosystem service provisioning. Although we here use the Barents Sea as a case study and provide insights in that particular system, our approach can be adapted to any type of ecological network.

## References

Artsdatabanken (2021). Rødlista 2021 - Artsdatabanken.

Beng, K.C. & Corlett, R.T. (2020). Applications of environmental dna (edna) in ecology and conservation: opportunities, challenges and prospects. Biodiversity and Conservation, 29, 2089–2121.

Cardinale, B.J., Duffy, J.E., Gonzalez, A., Hooper, D.U., Perrings, C., Venail, P., Narwani, A., Mace, G.M., Tilman, D., Wardle, D.A. et al. (2012). Biodiversity loss and its impact on humanity. Nature, 486, 59–67.

Cirtwill, A.R., Eklöf, A., Roslin, T., Wootton, K. & Gravel, D. (2019). A quantitative framework for investigating the reliability of empirical network construction. Methods in Ecology and Evolution, 10, 902–911.

Curtsdotter, A., Binzer, A., Brose, U., de Castro, F., Ebenman, B., Eklöf, A., Riede, J.O., Thierry, A. & Rall, B.C. (2011). Robustness to secondary extinctions: comparing trait-based sequential deletions in static and dynamic food webs. Basic and Applied Ecology, 12, 571–580.

Davies, T.W., Jenkins, S.R., Kingham, R., Kenworthy, J., Hawkins, S.J. & Hiddink, J.G. (2011). Dominance, biomass and extinction resistance determine the consequences of biodiversity loss for multiple coastal ecosystem processes. PloS one, 6, e28362.

Dee, L.E., Allesina, S., Bonn, A., Eklöf, A., Gaines, S.D., Hines, J., Jacob, U., McDonald-Madden, E., Possingham, H., Schröter, M. et al. (2017a). Operationalizing network theory for ecosystem service assessments. Trends in ecology & evolution, 32, 118–130.

Dee, L.E., De Lara, M., Costello, C. & Gaines, S.D. (2017b). To what extent can ecosystem services motivate protecting biodiversity? Ecology letters, 20, 935–946.

Díaz, S.M., Settele, J., Brondízio, E., Ngo, H., Gueze, M., Agard, J., Arneth, A., Balvanera, P., Brauman, K., Butchart, S. et al. (2019). The global assessment report on biodiversity and ecosystem services: Summary for policy makers.

Dunne, J.A., Williams, R.J. & Martinez, N.D. (2002). Network structure and biodiversity loss in food webs: robustness increases with connectance. Ecology letters, 5, 558–567.

Eklöf, A. & Ebenman, B. (2006). Species loss and secondary extinctions in simple and complex model communities. Journal of animal ecology, 75, 239–246.

Eklöf, A., Tang, S. & Allesina, S. (2013). Secondary extinctions in food webs: a Bayesian network approach. Methods in Ecology and Evolution, 4, 760–770.

Fossheim, M., Primicerio, R., Johannesen, E., Ingvaldsen, R.B., Aschan, M.M. & Dolgov, A.V. (2015). Recent warming leads to a rapid borealization of fish communities in the Arctic. Nature Climate Change.

Hansen, C., Skern-Mauritzen, M., van der Meeren, G.I., Jahkel, A. & Drinkwater, K. (2016). Set-up of the Nordic and Barents Seas (NoBa) Atlantis model. Fisken og havet nr. 2, p. 110.

Häussler, J., Barabás, G. & Eklöf, A. (2020). A bayesian network approach to trophic meta-communities shows that habitat loss accelerates top species extinctions. Ecology Letters, 23, 1849–1861.

HELCOM (2009). Biodiversity in the Baltic Sea ? An integrated thematic assessment on biodiversity and nature conservation in the Baltic Sea. Baltic Sea Environment Proceedings, 116B.

Holma, M., Laurila, E., Ahtiainen, H., Hallberg, M., Kull, M., Lepland, T., Oinonen, S. & Urb, J. (2019). Assessing economic, social, cultural and ecosystem services impacts in maritime spatial planning (MSP) in the Baltic Sea region.. Pan Baltic Scope 2019.

ICES (2021). Barents Sea Ecoregion – Ecosystem overview. report, ICES Advice: Ecosystem Overviews.

IUCN (2023a). The IUCN Red List of Threatened Species.

IUCN (2023b). Threats Classification Scheme (Version 3.3).

Jacob, U., Beckerman, A.P., Antonijevic, M., Dee, L.E., Eklöf, A., Possingham, H.P., Thompson, R., Webb, T.J. & Halpern, B.S. (2020). Marine conservation: towards a multi-layered network approach. Philosophical Transactions of the Royal Society B, 375, 20190459.

Keyes, A.A., McLaughlin, J.P., Barner, A.K. & Dee, L.E. (2021). An ecological network approach to predict ecosystem service vulnerability to species losses. Nature Communications, 12, 1–11.

Kivelä, M., Arenas, A., Barthelemy, M., Gleeson, J.P., Moreno, Y. & Porter, M.A. (2014). Multilayer networks. Journal of complex networks, 2, 203–271.

Kortsch, S., Primicerio, R., Aschan, M., Lind, S., Dolgov, A.V. & Planque, B. (2018). Food-web structure varies along environmental gradients in a high-latitude marine ecosystem. Ecography, pp. 1–14.

Kortsch, S., Primicerio, R., Aschan, M., Lind, S., Dolgov, A.V. & Planque, B. (2019). Food-web structure varies along environmental gradients in a high-latitude marine ecosystem. Ecography, 42, 295–308. eprint: https://onlinelibrary.wiley.com/doi/pdf/10.1111/ecog.03443.

Lyons, K.G., Brigham, C., Traut, B. & Schwartz, M.W. (2005). Rare species and ecosystem functioning. Conservation biology, 19, 1019–1024.

Maxwell, S.L., Fuller, R.A., Brooks, T.M. & Watson, J.E. (2016). Biodiversity: The ravages of guns, nets and bulldozers. Nature News, 536, 143.

McBride, M.M., Hansen, J.R., Korneev, O. & Titov, O. (2016). Joint Norwegian - Russian environmental status 2013. Report on the Barents Sea Ecosystem. Part II - Complete report. Tech. Rep. 2.

McComb, B.C. & Cushman, S.A. (2020). Synergistic effects of pervasive stressors on ecosystems and biodiversity. Frontiers in Ecology and Evolution, 8, 398.

McDonald-Madden, E., Sabbadin, R., Game, E.T., Baxter, P.W., Chades, I. & Possingham, H.P. (2016). Using food-web theory to conserve ecosystems. Nature communications, 7, 1–8.

OSPAR (2024). List of Threatened and/or Declining Species & Habitats.

Pilosof, S., Porter, M.A., Pascual, M. & Kéfi, S. (2017). The multilayer nature of ecological networks. Nature Ecology & Evolution, 1, 1–9.

Planque, B., Primicerio, R., Michalsen, K., Aschan, M., Certain, G., Dalpadado, P., Gjøsæater, H., Hansen, C., Johannesen, E., Jørgensen, L.L., Kolsum, I., Kortsch, S., Leclerc, L.M., Omli, L., Skern-Mauritzen, M. & Wiedmann, M. (2014). Who eats whom in the Barents Sea: A food web topology from plankton to whales. Ecology, 95, 1430–1430.

Poisot, T., Mouquet, N. & Gravel, D. (2013). Trophic complementarity drives the biodiversity– ecosystem functioning relationship in food webs. Ecology letters, 16, 853–861.

Poisot, T., Stouffer, D.B. & Gravel, D. (2015). Beyond species: why ecological interaction networks vary through space and time. Oikos, 124, 243–251.

Ryser, R., Häussler, J., Stark, M., Brose, U., Rall, B.C. & Guill, C. (2019). The biggest losers: habitat isolation deconstructs complex food webs from top to bottom. Proceedings of the royal society B, 286, 20191177.

Smith, M.D. & Knapp, A.K. (2003). Dominant species maintain ecosystem function with non-random species loss. Ecology Letters, 6, 509–517.

Staniczenko, P.P., Sivasubramaniam, P., Suttle, K.B. & Pearson, R.G. (2017). Linking macroe-cology and community ecology: refining predictions of species distributions using biotic interaction networks. Ecology letters, 20, 693–707.

Tilman, D., Clark, M., Williams, D.R., Kimmel, K., Polasky, S. & Packer, C. (2017). Future threats to biodiversity and pathways to their prevention. Nature, 546, 73–81.

Winfree, R. W. Fox, J., Williams, N.M., Reilly, J.R. & Cariveau, D.P. (2015). Abundance of common species, not species richness, drives delivery of a real-world ecosystem service. Ecology letters, 18, 626–635.

Xiao, H., Dee, L.E., Chades, I., Peyrard, N., Sabbadin, R., Stringer, M. & McDonald-Madden, E. (2018). Win-wins for biodiversity and ecosystem service conservation depend on the trophic levels of the species providing services. Journal of Applied Ecology, 55, 2160–2170.

