## Supplementary information for "Cascading effects of species loss on ecosystem service provisioning in a marine food web"

[†] Corresponding author

#### 1 Description of threats

##### 1.1 Selective extraction of species

"Selective extraction of species" refers to the removal or intentional harvest of particular species, including both target and incidental catch.

The majority of all landings in the Barents Sea are made with demersal (bottom) trawlers, but also longlines, gillnets, handlines, floatlines, bottom seiners, Danish seiners, purse-seiners, pelagic trawlers, and crab pots (ICES (2021b)).

The main **pelagic** fish species targeted by commercial fisheries in the Barents Sea is capelin, *Mallotus villosus*. The main targeted **demersal** species are cod, *Gadus morhua*, haddock, *Melanogrammus aeglefinus*, saithe, *Pollachius virens*, and other gadoids (including polar cod, *Boreogadus saida*). The main targeted crustaceans are deep-sea prawn, *Pandalus borealis*, red king crab, *Paralithodes camtchaticus*, and snow crab, *Chionoecetes opilio*. Targeted mammals are harp seals, *Pagophilus groenlandicus*, and minke whales, *Balaenoptera acutorostrata*. Additionally, there are also minimal catches of herring, *Clupea harengus*, mackerel, *Scomber scombrus*, and blue whiting, *Micromesistius poutassou* (ICES (2021b)).

The largest populations of commercially fished species are capelin, cod and haddock; all three are assessed to have full reproductive capacity (ICES (2021b)). The capelin population fluctuates a lot naturally as it is a short-lived species and population size therefore depends on recruitment success (Bogstad *et al.* (2021); Hesthagen *et al.* (2021)). The last four decades the population has collapsed three times (1985-1989, 1993-1997, and 2003-2006) after periods of intense predation pressure and intense fishing (landings of up to 3 million tonnes). After each collapse there have been years with total fishing bans on capelin, and today the management plan for capelin is quite restrictive (ICES (2021b)). Capelin is assessed as LC (Least concern) by Artsdatabanken (Hesthagen *et al.* (2021), but its population size continues to fluctuate a lot between years (Bogstad *et al.* (2021)).

Two fish populations are relatively small and are assessed to be overfished: golden redfish, *Sebastes norvegicus* and Norwegian coastal cod (ICES (2022); ICES (2021b)). *Sebastes spp.* are particularly sensitive to over exploitation because they have slow growth and mature late (Christiansen & Reist (2013)).

In 2021, ICES provided fishery advice for 16 species in the Barents Sea. These include the most important commercial species, but some species that are fished to a lesser extent lack advice, including the ecologically important polar cod, *Boreogadus saida*, which is mainly fished by Russia (Christiansen & Reist (2013)). Some other unassessed populations that are fished in the BS are wolffishes, *Anarhichadidae spp.*, plaice, *Pleuronectes platessa*, and anglerfish, *Lophius spp.* (ICES (2021b)).

Even when a population is not considered overfished, all harvesting amplifies variability in fish populations, which destabilizes them (Christiansen & Reist (2013)).

An unavoidable consequence of targeted commercial fishing is that non-target species will get caught in the fishing gear as *bycatch*. This poses a significant threat to some groups of species, such as seabirds. In this section, and when linking threats to species in the model, catches in active fishing gear is considered bycatch while catches/entanglement in lost/discarded fishing gear is considered litter (see Litter) but the effects on animals are similar.

Population impacts of bycatch on seabirds in the Barents Sea are largely unknown because of lacking data (ICES (2021b)), but globally we know that bycatch is one of the top threats to seabirds (Dias *et al.* (2019)). Birds are caught by trawls, longlines and gillnets, and especially the use of nylon fishnets and longlines in fisheries has greatly increased the number of seabirds caught in fishing gear (Furness (2003)). Coastal and pelagic diving seabirds, such as *Alcidae* species (aulks), are more frequently caught by gillnets while surface-feeding seabirds, such as *Fulmarus glacialis* (northern fulmar), are more often caught by longlines (ICES (2021b)). *F. glacialis* is one of the bird species caught in the highest numbers in longline-fisheries globally (Anderson *et al.* (2011)), but the population effects are unknown. Both Žydelis *et al.* (2009) and Anderson *et al.* (2011) state that records of seabird bycatch are most likely underestimated for both gillnet and longline fisheries.

Other than seabirds, non-target fish species, elasmobranchs, and harbour porpoises are caught as bycatch in the Barents Sea. The data is lacking on frequencies and population impacts of bycatch on elasmobranchs, especially sharks, but it is known that skates are affected and that populations of large-bodied elasmobranchs could potentially be extra vulnerable. It is estimated that about 7 000 harbour porpoises, *Phocoena phocoena*, are caught in gillnets every year, but the population consequences of this are unknown (ICES (2021b)). Fish species with benthic and resident lifestyles and long generation times are more vulnerable to bycatch (Christiansen & Reist (2013)).

The main regulation of the Barents Sea fisheries are the TACs (Total Allowable Catch) that are decided by the Joint Norwegian-Russian Fisheries Commission. Coastal fisheries of Russia and Norway are managed by the respective national authorities. The Joint Norwegian-Russian Fisheries Commission also manages seal hunting, while quotas for minke whale hunting are set by Norwegian authorities based on the International Whaling Committee's (IWC) Revised Management Procedure (RMP) (ICES (2021b)).

For the threat scorings of "Selective extraction of species", the following literature was used:

- Fisheries Overview (ICES (2022))
- Joint Norwegian - Russian environmental status 2013. Report on the Barents Sea Ecosystem. Part II - Complete report (McBride *et al.* (2016))
- Red List assessments from the Norwegian 2021 Red List (Artsdatabanken (2021))

In 2022 ICES did not assess cod, haddock, capelin, prawn, and beaked redfish because of the "temporary suspension of the Russian Federation from ICES activities" (ICES (2022)). Therefore, the 2021 assessment (ICES (2021b)) has been used for those species.

#### 1.2 Ocean acidification

When atmospheric $CO_2$ dissolves in water, the net effects are higher concentrations of $H_2CO_3$, $HCO_3^-$, and $H^+$ and lower concentrations of $CO_3^{2-}$ (carbonate ion). Since pH = -log[$H^+$], the result is a decrease in pH (acidification). The decrease in [$CO_3^{2-}$] means that there is less aragonite and calcite (the two forms of $CaCO_3$; calcium carbonate) available for calcifying organisms to use (Fabry *et al.*, 2008).

Studies have shown that the sensitivity of some species to ocean acidification, at least partly, depends on available energy and protective mechanisms. Some species seem to be more resistant to acidification, and can calcify even in undersaturated waters, if they have large enough food resources (AMAP, 2013). Resistance to acidification also depends on water temperature; increased temperature can amplify the negative effects of acidification—giving cause for concern for future tandem effects under global warming (Rodolfo-Metalpa *et al.*, 2011).

The Arctic Ocean and its marginal seas (including Barents Sea) is one of the regions most sensitive to ocean acidification, because it is naturally poorly buffered and has low concentrations of $CO_3^{2-}$ (AMAP, 2013; Fabry *et al.*, 2008). The northern polar seas have already shown seasonal under-saturation of aragonite and are projected to be under-saturated year-round by the mid-21st century.

The rapid acidification of the Arctic Ocean is being driven by increasing levels of atmospheric $CO_2$ from anthropogenic sources, and is further spurred on by melting sea ice and increasing carbon input from terrestrial sources (Qi et al., 2022). Melting ice worsens the under-saturation of aragonite and calcite in two ways: 1) larger areas of open water leads to more $CO_2$ moving from the atmosphere into the water, and 2) the ice consists of brackish water with low concentrations of $Ca^{2+}$, so the meltwater decreases [$Ca^{2+}$] of the whole water mass (AMAP, 2013).

There are multiple other reasons why acidification could have a particularly large impact in the Arctic Ocean region. Since these ecosystems are highly productive, a lot of $CO_2$ is already produced, and the Barents Sea is no exception — having one of the most productive ecosystems at this latitude (Carmack & Wassmann, 2006). There is also potential for increasing inputs of carbon from other sources; large amounts of DOC and POC (dissolved and particulate organic carbon, respectively) are flowing from rivers and eroding coasts into the Arctic shelf seas. Global warming also causes more methane ($CH_4$) to be released from sediments into the water, where it contributes to acidification by metabolising into $CO_2$ and to oxygen depletion by oxidising (AMAP, 2013).

The pH and aragonite saturation values in the Barents Sea vary both seasonally and geographically because of the large variability in the inflow waters. There is also a lack of data, especially from the northern and eastern Barents Sea. Therefore, it has been difficult to assess trends in ocean acidification (AMAP, 2018). By 2013, calcium carbonate undersaturation had not been observed in the Barents Sea, but it was projected to partially reach aragonite undersaturation in the near future, based on measured changes in bottom waters (AMAP, 2013). There is a varying but overall decreasing pH and calcium carbonate saturation, which indicates increasing acidification levels in the Barents Sea (Freitas *et al.*, 2022; Fransner *et al.*, 2022).

Skogen *et al.* (2014) projected that the surface water pH in the Barents Sea will decrease by 0.19 units on average, and up to 0.25 units in the northern region, by 2065 (quite a large decrease). Additionally, Wallhead *et al.* (2017) projected that the bottom pH in the Barents Sea would decrease by 0.1-0.2 units in the coming 50 years, and that the bottom waters would experience aragonite undersaturation by 2070.

Organism groups in the Barents Sea that are especially sensitive to ocean acidification are mollusks, especially thecosomatous pteropods, such as *Limacina sp.*, and other thin-shelled planktonic mollusks, foraminifera, and echinoderms. Another affected group is crustaceans, such as crabs of the genus *Hyas*, barnacles *Balanus sp.*, and potentially *Calanus* copepods (AMAP,

2013; Fabry *et al.*, 2008). Studies have also shown detrimental effects of acidification on the fishes *Boregodaus saida* and *Gadus morhua* (AMAP, 2018). For all groups, studies have shown varying effects on survival depending on species, life stage, region, and study design, and more research is needed (AMAP, 2018).

#### 1.3 Oil pollution

The overall level of oil pollution in the Barents Sea is relatively low compared to other European marginal and inland seas. There are internal differences in oil pollution between regions though: the southern region, where there is most ship traffic, has the most oil pollution and the central region, where there are most fisheries, has a medium amount of pollution— while the northern region is less polluted (Ivanov *et al.*, 2022). Arctic ecosystems are possibly more vulnerable to oil spills than other marine ecosystems; Arctic winter conditions slow down the degradation and loss of hydrocarbons and the logistics of cleaning up spills are complicated by the ice, leading to longer exposure of fauna (AMAP, 2010; Barry *et al.*, 2013).

The risk of oil spill accidents in the Barents Sea has increased in the last decade and will increase further in the coming one, due to increasing hydrocarbon activities and ship traffic (ICES, 2019). Multiple small spills and chronic exposure can impact the ecosystem as much as, or more than, one large spill (AMAP, 2010).

Population effects of oil pollution are often hard to measure, and for many organisms hard or impossible to study experimentally. Some habitats, macroalgae, and invertebrates can also be damaged by clean-up operations following spills and the effects can be hard to disentangle (AMAP, 2010). Additionally, few studies have focused on Arctic species or the Barents Sea.

An important, and complicating, aspect regarding population effects of oil pollution is that they will vary significantly depending on the timing and location of a spill. For example, even a moderately sized oil spill in a nesting area during the nesting period, when tens of thousands of birds are confined to the same area, could have catastrophic consequences for a bird population. Another example that could have large detrimental effects is an oil spill occurring near the ice during the polar and Arctic cod (*Boreogadus saida* and *Arctogadus glacialis*) spawning and/or egg incubation period (AMAP, 2010).

In general, seabirds and mammals with fur are highly sensitive to oil spills; oil from the sea surface can cover their pelt or plumage (oiling) where it destroys its insulating properties, resulting in hypothermia and often death. Heavy oiling can also hinder seal pups from swimming, killing them (AMAP, 2010). Birds that during some part of the year swim and/or feed at sea (e.g., Little auk, *Alle alle*) are extra vulnerable to oil spills (Fort et al., 2013). Incubating birdeggs are also very sensitive and even light oiling can cause death or mutations (AMAP, 2010). Out of all Arctic birds, alcids and fulmars are most seriously affected by acute oil spills because their population growth is mostly governed by adult survival (AMAP, 2010).

Fishes that have long egg incubation times, such as the flatfish *Hippoglossus hippoglossus*, and/or spawn eggs under sea ice, such as *Boreogadus saida*, could be more vulnerable to oil spills (AMAP, 2010).

In the AMAP report AMAP Assessment 2007: Oil and gas activities in the Arctic - effects and potential effects (AMAP, 2010), the authors stated that since the Arctic has generally low concentrations of oil hydrocarbons and PAHs, adverse effects on fauna are not expected other than in locally contaminated areas. While oil-related PAH concentrations in the Barents Sea are generally low by international standards, multiple lines of evidence indicate that some Arctic PAH metrics have not declined—and can rise under recent climate and local-emission influences (Boitsov *et al.*, 2009; Yu *et al.*, 2019). In addition, the risk of oil spills has increased since the AMAP assessment was written and will continue to increase (ICES, 2019). Additionally, considering that oil pollution levels vary regionally in the Barents Sea (Ivanov *et al.*, 2022), it can not be assumed today that no fauna will experience adverse effects from chronic oil pollution. These effects and their extent are, however, not well known.

##### 1.4 Increased water temperature and ice loss

Global warming is causing the Arctic to experience ice loss and increasing sea surface temperatures (SSTs). Warming rates in the Arctic are currently two to four times higher than the global average ((Rantanen *et al.*, 2022) — a phenomenon called Arctic amplification (Serreze & Barry, 2011).

Water temperatures in the Barents Sea can naturally fluctuate quite quickly, largely depending on the amount of inflow from the Atlantic, but the overall temperature has been rising during the last four decades (Watelet *et al.*, 2020; Skagseth *et al.*, 2020). The northern Barents Sea is the Arctic area with the highest rates of surface warming and sea ice loss (Screen & Simmonds (2010)) and it is expected to become completely ice-free year-round before the end of this century (Onarheim & Årthun (2017)). Imported sea ice from the interior Arctic controls salinity levels and keeps the northern Barents Sea cold and stratified, but with less imported sea ice the northern Barents Sea water column is now shifting towards Atlantic water-properties: less saline, warmer, and less stratified (Lind *et al.* (2018)). This so-called *Atlantification* of the northern Barents Sea is causing a *borealisation* of the ecosystem, where Atlantic species expand northwards while Arctic species decline (Fossheim *et al.* (2015)).

Primary production from pelagic phytoplankton and ice-associated (sympagic) algae supports Arctic food webs from the bottom up. Ice algae is especially crucial for spring production as it needs less light than pelagic phytoplankton to bloom, but it has also been shown to be an important carbon source year-round (Koch *et al.*, 2023). The importance of ice algae in supporting the food web is further illustrated by the fact that the keystone secondary producer *Calanus glacialis* times its reproduction to the spring ice algae blooms (Leu *et al.*, 2011). In the Arctic as a whole, climate change has temporarily created more suitable habitat for ice algae, with thinner ice and less snow cover, but in the Barents Sea the overall decrease in ice cover already outweighs the increase in habitat suitability (Lim *et al.*, 2022). The loss of sea ice also results in more light influx to the water column, increasing the production of pelagic phytoplankton (Horvat *et al.*, 2017). A decrease of ice algae and increase of pelagic phytoplankton will have consequences for the whole food web (Dalpadado *et al.*, 2020).

Climate change is already affecting zooplankton population composition: lipid rich euphausiids have increased in numbers while Arctic species such as *Calanus glacialis* and *Themisto libellula* have decreased (Dalpadado *et al.*, 2012). (Dalpadado *et al.*, 2012) write that a possible future scenario is that *C. glacialis* and *C. hyperboreus* will decrease in favour of *Calanus finmarchicus*, as the Barents Sea becomes more Atlantified and *C. finmarchicus* from the North Sea migrates to the Barents Sea.

The population of the keystone species polar cod *Boreogadus saida* has seen a large decline and migration northwards in the last 10 years (Gjøsæter *et al.*, 2020; Hesthagen *et al.*, 2021) Warming waters are likely hindering recruitment of the polar cod, as well as the less abundant ice cod (*Arctogadus glacialis.* These fishes are endemic to the Arctic and highly adapted to Arctic water conditions, preferring temperatures between 2–5.5°C (Gjøsæter *et al.*, 2020; Hesthagen *et al.*, 2021). Polar cod and ice cod are also cryopelagic, meaning that they use sea ice as habitat and spawning substrate (Huserbråten *et al.*, 2019). Declines in the polar cod population is likely to initiate a regime shift in the marine Arctic (Christiansen & Reist, 2013).

Marine mammals such as *Ursus maritimus* and several phocid species are obligatorily—or at least partially—dependent on sea-ice platforms for reproductive activities and foraging (Bajzak *et al.*, 2011; Johnston *et al.*, 2012; Rode *et al.*, 2021; Archer *et al.*, 2025). The progressive loss of sea ice therefore eliminates critical habitat, breeding sites, and/or hunting arenas for these taxa. Long-term monitoring of the Svalbard polar-bear sub-population indicates a pronounced demographic decline over the past several decades (Vongraven *et al.*, 2023), and physiological assessments reveal that a substantial fraction of the extant individuals now display markedly reduced body-condition indices (Archer *et al.*, 2025).

The organisms forming important Barents Sea habitats such as coral reefs, coral gardens, and kelp forests are also thought to be negatively affected by warming, since they have relatively low temperature optima (OSPAR, 2022a).

An indirect effect of the occurring ice loss is that it increases the accessibility of the Barents Sea to activities that cause other threats to the ecosystem: ship traffic, fishing, hydrocarbon exploration, and migrating organisms (Smith & Stephenson, 2013). Increased migration and changed migration patterns of animals into the Arctic due to sea-ice reductions could lead to an increase in viral diseases such as PDV (see *Pathogens*) (VanWormer *et al.*, 2019). Migration and transport of invasive alien species can also lead to larger problems with these species.

Ice melting is also expected to increase the levels of plastic pollution in the whole Arctic Ocean, since sea ice contains large amounts of macro- and microplastics (Collard & Ask, 2021) (see *Marine litter*).

#### 1.5 Habitat destruction

While the threat "Seabed abrasion" covers the direct damage of trawling gear to organisms, "Habitat destruction" here refers to the indirect impacts on organisms when their habitats are damaged/destroyed. This excludes ice habitats which are treated under "Ice loss".

The"OSPAR list of Threatened and/or Declining species and habitat" (OSPAR (2024)) lists eight habitats that are present in the Barents Sea:

- **Coral gardens**
- **Deep-sea sponge aggregations**
- **Kelp forests**
- **Lophelia pertusa reefs**
- ***Modiolus modiolus* beds**
- **Seamounts**
- Sea pen and burrowing megafauna communities
- (Carbonate mounds)

Habitats we have linked to taxa present in the Barents Sea food web dataset are listed in **bold** and described below.

**Lophelia pertusa reefs**

Reefs of the hard coral *Lophelia pertusa* occur on hard substrates, mainly in water temperatures in the interval 4-8°C (OSPAR (2009)). The reefs make up structurally complex habitats which support a high biodiversity of associated fauna (OSPAR (2009); Buhl-Mortensen (2017)).

In the Barents Sea, associated fauna includes sponges, Norway redfish *Sebastes viviparus*, tusk *Brosme brosme*, saithe *Pollachius virens*, and soft corals of the genus *Nephtheidae* (Buhl-Mortensen, 2017; Kutti *et al.*, 2015).

*Lophelia pertusa* reefs are especially threatened by seabed abrasion from trawling gear. They are also negatively impacted by pollution, siltation (from dumping of waste and dredged material), eutrophication, and to some extent damage from tourism (diving) (OSPAR (2009)). They are also threatened by acidification caused by climate change (AMAP (2013); OSPAR (2022b)). Living *L. pertusa* corals seem to be quite resistant to acidification, but dead corals –which make up large parts of the coral reefs– are more vulnerable to dissolution under acidification. Therefore, acidification threatens these habitats by causing the reefs to collapse (AMAP (2018)).

**Coral gardens**

Coral gardens are aggregations of mostly non-reef forming corals, such as scleractinian (hard) corals, leather corals, Alcyonacea (represented by genus *Nephtheidae* in the BS food web dataset), and anemones, *Actiniaria sp.*. They can also have some reef-forming corals such as *Lophelia pertusa*, but not in majority (OSPAR (2008a)).

While there is little research on coral garden communities in the Barents Sea, Krieger & Wing (2002) found that, in Alaska, species such as *Sebastes* fishes, shrimps, sea anemones, basket stars, and crabs were associated with coral gardens.

Coral gardens are threatened by trawling and long-lining that causes damage to and physical removal of species. Climate change induced ocean acidification is also a threat, especially to aragonite corals (see 5.1.2 Ocean acidification). Other potential threats are increased sea temperature, marine litter, oil and chemical pollution, deep-sea mining, and aquaculture waste (OSPAR (2022b)).

**Deep-sea sponge aggregations**

Sponges are filter-feeding invertebrates in the order Porifera. The sponges making up deep-sea sponge grounds in the Barents Sea belong to the classes Hexactinellida and Demospongiae (represented by genus *Geodia* in the BS food web dataset). Sponges prefer similar habitats as cold-water corals (*L. pertusa* reefs) and they can often be found in the same areas (OSPAR (2008b)). Dense sponge aggregations are common in the Barents Sea(Jørgensen *et al.* (2016)). They create structurally unique habitats because their spicules aggregate around the sponge grounds, preventing burrowing fauna from inhabiting the area (OSPAR (2008b)).

Sponge grounds support high species richness and are important habitat for many species (Jørgensen *et al.* (2016)); they are especially ideal habitat for brittle stars (order Ophiurida) (OSPAR (2008b)). Some fishes, such as *Sebastes viviparus* and *B. brosme*, have similar affinities for sponge grounds and other structurally complex habitats, while others, such as *Chimaera monstrosa*, have been found to prefer sponge grounds (Kutti *et al.* (2015)).

Deep-sea sponge aggregations are mainly threatened by trawling damage and changes in temperature, pH, and circulation caused by climate change (OSPAR (2022b)).

**Kelp forests**

Kelp forests are coastal habitats made up of brown macroalgae; in the Barents Sea the forest-forming species are mainly *Laminaria hyperborea* and *Saccharina latissima* (OSPAR (2021b)).

Kelp forests are important habitats for epiphytic algae, epiphytic fauna, such as tunicates, sponges, and bryozoans, and invertebrates such as snails, bivalves, crustaceans, and polychaetes. Many fish species, such as the commercially important cod and pollack also use kelp forests as nurseries and feeding grounds (Gundersen *et al.* (2017)).

For the last five decades, kelp forests in the BS region have been overgrazed by sea urchins (*Strongylocentrotus droebachiensis*) to the point where they have mostly become barren grounds. This development has however turned around slightly in the last two decades and the kelp forests are recovering (Gundersen *et al.* (2017)).

Kelp forests are also sensitive to increases in water temperature because the kelp species are adapted to cold temperatures. Other threats to kelp forests, that are less prominent in the Barents Sea region, are pollution, storms (lead to higher turbidity which hinders photosynthesis), eutrophication (limitates photosynthesis through excessive growth of kelp epiphytes and phytoplankton), and seabed abrasion (causes physical damage and higher turbidity) (OSPAR (2021b); OSPAR (2021a); Gundersen *et al.* (2017)).

**_Modiolus modiolus_ beds**

Horsemussel, *M. modiolus*, beds are abundant in the Barents Sea, although not well surveyed or studied. *M. modiolus* beds are not especially fragile but they are sensitive to abrasion and other physical impact. They recover very slowly from damage due to their slow and sporadic recruitment. In other regions, *M. modiolus* beds are known to support a large invertebrate community. Examples of associated taxa are *Hyas spp.*, *Balanus spp.*, solitary ascidians, and sponges (OSPAR (2008c)).

**Seamounts**

Seamounts are underwater mountains, usually inactive volcanoes. They serve as substrate for fauna and provide ideal conditions for suspension feeders such as sponges, hydroids, and ascidians. Seamounts are in general not very well studied. Because of their structural stability, seamounts are not currently threatened, but their communities are impacted by trawling, and the mounts could be threatened in the future by extraction of mineral resources (OSPAR (2010); OSPAR (2022b)).

#### 1.6 Marine litter

Marine litter is defined as "any persistent, manufactured or processed solid material discarded, disposed of or abandoned in the marine and coastal environment" Werner & O'Brien (2017) . Marine litter occurs in all of the world's oceans. Marine litter has both ocean-based sources, mainly maritime transport, fisheries, aquaculture, and oil and gas platforms, and land-based sources, mainly landfills, untreated sewage water, industries, medical facilities, and tourism.

The Arctic Ocean has a disproportionately large amount of litter considering its area and the low population densities of surrounding land areas. This is because of atmospheric and oceanic movements, large river inflows from land, and local ocean-based activities (Bergmann *et al.*, 2022). In the Barents Sea, the local ocean-based sources of litter include fisheries, maritime transportation, hydrocarbon exploration, tourism, and aquaculture. As expected, local land-based activities represent a smaller proportion of the litter input (Bergmann *et al.*, 2022; Grøsvik *et al.*, 2018).

In the joint Norwegian-Russian ecosystem monitoring survey running from 2010 to 2016, Grøsvik *et al.* (2018) found marine litter in all parts of the Barents Sea with plastic being the dominating material in the pelagic region and wood dominating in the surface and benthic region. They also found indications of there being more plastic litter in areas with more traffic from transport and fisheries.

Marine litter can be of any material but plastic is particularly problematic and widespread; large plastics can smother benthic habitats and entangle animals of all sizes, and as they get broken down into smaller fragments (secondary micro- and nanoplastics), through abiotic processes and animal consumption, the plastic becomes available to organisms of lower and lower trophic levels (Kühn *et al.*, 2015; Zhang *et al.*, 2025). In the Barents Sea, abandoned fishing gear is an especially large source of plastic litter and microplastics, compared to other regions (Bergmann *et al.*, 2022).

There are three main ways that marine litter impacts wildlife negatively:

- **Entanglement.** Animals can get stuck in litter such as large plastics (bags, balloons, synthetic ropes, etc.) and lost fishing gear (nets and traps) (ghost fishing). The entangled animal can get directly hurt or suffocated by the litter, and/or get hindered in their behavior and can thus starve or drown.

  Any animal species can become entangled in litter and although the long-term effects of entanglement on populations are usually hard to estimate, it has been shown to negatively impact the survival of some populations (Kühn *et al.*, 2015).

- **Ingestion.** As with entanglement, species of all trophic levels can be affected by ingestion of marine litter, either firsthand or secondhand (through consumption of prey which has ingested litter).

  Ingested litter can accumulate in the stomach which limits the individual's food intake and can reduce the efficiency of digestion, leading to poor nutrition, dehydration, and overall worsened body condition. It can also block and/or damage the gastrointestinal tract which can shorten the individual's life-span or, in the worst case, lead to a rapid death.

  As stated above, ingestion can cause direct mortality but it probably does not do so at a frequency that is relevant at the population level. Rather, the sub-lethal decreases in body condition is what lowers average survival and reproduction rates at the population level (Kühn *et al.*, 2015).

- **Smothering.** Larger debris can cover, damage, and suffocate (cause anoxic conditions in) the seabed and its habitants (e.g., corals, sponges, and seagrasses). This has both direct and indirect effects on the food web (Kühn *et al.*, 2015).

Outside of these main effects, synthetic materials can also be used as new habitats and act as vectors for migration, potentially facilitating spread of invasive alien species (Kühn *et al.*, 2015).

The effects of marine litter on populations can be hard to translate into threat impact scores, but since litter can impact organisms through multiple pathways and is ever present it is at least safe to assume that this threat impacts whole populations of most species (Zhang *et al.*, 2025). As Kühn *et al.* (2015) writes: "... given sufficient time and research effort, all species of marine organisms will get documented examples of interaction with marine debris".

#### 1.7 Seabed abrasion

Seabed abrasion is perhaps one the most publicly well-known threats to marine environments world-wide. It is the physical removal or hurting of benthos and it is mainly caused by trawling. As such, it is directly linked to commercial fishing (ICES, 2021a). Seabed abrasion caused by trawling removes important habitat-forming species such as sponges, kelp, and hard and soft corals, as well as other benthic fauna. This has both direct and indirect effects on the ecosystem (OSPAR, 2009, 2008b). Here,"Seabed abrasion" refers to direct impacts of abrasion to organisms, since habitat destruction is treated as its own threat.

Density and diversity of megabenthos (benthic organisms ¿ 2cm) in the Barents Sea has been shown to be negatively correlated with trawling intensity (Buhl-Mortensen *et al.*, 2016; Jørgensen *et al.*, 2016). Traits that make a species particularly vulnerable to being caught in a trawl are being large, high above the sediment, and immobile. Using the former two traits as criteria, Jørgensen *et al.* (2016)) identified 23 high-risk megabenthic species and 80 medium-risk species in the Barents Sea, and showed that the most vulnerable species had the strongest negative relationships between biomass and trawling. Some of the identified high-risk species are basket stars, *Gorgonocephalus sp.*, Geodia sponges, and sea cucumber, *Cucumaria frondosa.*

Bottom trawling also increases ocean acidification and global warming by remineralizing sedimentary carbon to aqueous $CO_2$ (Sala *et al.*, 2021), of which, according to Atwood et al.(2024) a majority is released to the atmosphere.

#### 1.8 Acoustic disturbances

Underwater noise is caused by ship traffic, icebreaking, seismic airguns (used to find oil and gas), drilling, dredging, pile driving, military sonar, and aircrafts. There are large knowledge gaps regarding how Arctic marine animals are impacted by acoustic disturbances. There are few studies focusing on fish, and while mammals are generally better researched, some mammals have not been studied at all. There is also a lack of data on population-level effects, no studies examine effects of chronic noise exposure, and few studies have been conducted in the European Arctic (Halliday *et al.*, 2020; de Jong *et al.*, 2020).

Regardless, we do know that most marine vertebrates use sound or hearing for important functions such as navigation, communication, reproduction, and predator-prey interactions. We also know that noise pollution can disturb all of these functions through sound-masking and induced behavioural changes, as well as cause acute physical damage, hearing-loss, and chronic stress (Arctic Council, 2009; de Jong et al., 2020). Arctic marine animals are also thought to be especially sensitive to impacts of acoustic disturbances because of the Arctic's long history with low levels of anthropogenic noise and ambient sound (Halliday *et al.*, 2020).

The research on effects of noise pollution has mainly been focused on marine mammals, and the current consensus seems to be that mammals, and especially cetaceans, are particularly impacted because anthropogenic noise disturbs their echolocation and communication (and thus their migration, foraging, and reproduction) (Moore *et al.*, 2012). Mammals with narrow ecological niches and high site fidelity (i.e., low dispersal ability), such as narwhals *Monodon monoceros*, are particularly sensitive to all disturbances, including noise pollution (Tervo *et al.*, 2023).

#### 1.9 Other pollution

A big problem with pollution in (especially marine) environments is bioaccumulation and biomagnification, e.g., that lipophilic pollutants such as OHCs (organohalogen compounds) and MeHg (methylmercury) accumulate in the tissues of biota and that the concentrations increase with trophic level (AMAP, 2018). Hence, top-level consumers can accumulate detrimental levels of heavy metals and POPs.

For Arctic mammals and birds, high levels of POPs and MeHg can have varying negative effects on vitamin physiology, endocrinology, reproduction, immune function, and neurology. Relatively little is known about the impacts of contaminants on fish, despite the fact that they can accumulate relatively high levels of OHCs and mercury (AMAP, 2018).

The Arctic, including the Barents Sea, has relatively low pollution levels compared to more industrialized regions in the world (AMAP, 2018; ICES, 2021b). Persistent organic pollutants (POPs) have not been widely used in the Arctic, but they are transported to the region through atmospheric transport, ocean currents, and river inputs (AMAP, 2018). The most critical pollution-pathway for the Barents Sea is this long-range transport of POPs and heavy metals (Hansen et al., 2016). Other pathways of contamination for the Barents Sea are natural processes, ship fuel emissions, accidental local releases, fresh-water runoff, and industrial activities related to extraction of petroleum products and marine transport (ICES, 2021b; McBride *et al.*, 2016).

Other than increasing mortality of organisms, pollution also negatively impacts the ecosystem service *Food provisioning* by reducing the safety and quality of fish. This could have major consequences both for the health of people living in the Barents Sea region and for the Norwegian fishing industry.

#### 1.10 Pathogens

Pathogens can affect populations of species by killing or reducing the fitness of individuals. Pathogens have always existed, they are themselves part of the ecosystem, and humans usually aren't the main cause of their spread, but our presence and activities can increase the spread of pathogens. Because multiyear ice in the Barents Sea is melting (see *Ice loss*) new pathways for species migration, and therefore also spread of pathogens, will open up (VanWormer *et al.*, 2019). Here are some known pathogens that impact species in the Barents Sea.

Morbilliviruses such as PDV (Phocine Distemper Virus) and CeMV (Cetacean Morbillivirus) weakens the immune systems of hosts and can cause mass mortalities. PDV epidemics causing mass mortalities have occurred multiple times in northern European seal populations; harbour seals *Phoca vitulina* seem to be the most susceptible to death by PDV (Barratclough *et al.*, 2023). In the Barents Sea, susceptible seal species are harp seals, *Pagophilus groenlandicus* (Duignan *et al.*, 2014), ringed seals *Pusa hispida*, and hooded seals *Cystophora cristata* (Barratclough *et al.*, 2023). CeMV occurs in porpoises, dolphins, narwhals, and fin whales (Barratclough *et al.*, 2023).

*Brucella spp.* are coccobacillus which are known to cause infertility and abortion in terrestrial and marine mammals. No reproductive or mortality issues have so far been reported in Arctic marine mammals, but *Brucella* has been found in polar bears, seals, walruses, and some whales (Barratclough *et al.*, 2023).

The highly contagious and often fatal virus Highly pathogenic avian influenza (HPAI), a subtype of *Influenza A* virus, perhaps better known as 'bird flu', has been detected in bird populations (e.g. pink-footed geese *Anser brachyrhynchus*) breeding in Svalbard (Madslien *et al.*, 2021). *Influenza A* also occurs in white whales, ringed seals (Canadian Arctic), and potentially narwhals and bowhead whales (Barratclough *et al.*, 2023).

Botulism, caused by the toxin produced by the bacterium *Clostridium botulinum*, is a potential threat for birds like herring gulls *Larus argentatus* (Neimanis *et al.*, 2007).

Protozoan parasites such as *Toxoplasma gondii* can affect marine mammals. *T. gondii* currently has a low risk of causing mortalities in the Arctic, but it has been found in walruses on Svalbard and narwhals, polar bears, ringed seals, and bearded seals in other parts of the Arctic (Barratclough *et al.*, 2023).

### 2 Quantifying the impact of threats on individual taxa

Different threats are can have more or less severe effects, which also varies depending on the species exposed. We here standardized method for determining the *magnitude* of the different threats analyzed. We may also call this threat impact.

The IUCN determines threat impact scores for different threats and species in the Red List (with the exception of an ongoing temporary pause since Red List 2022-2) using a scoring scheme. Although the methods used to estimate species extinctions have been criticized and improvements to the criteria are widely sought after (Edgar, 2025; Zhao *et al.*, 2025), we base our scoring on the IUCN system. The scheme includes the factors timing, scope, and severity which each gets a scoring from 0 to 3; the scores are added together and the highest impact score a threat can get is then 9 (IUCN (2023)). IUCN's scheme is the basis of the scoring matrix (Fig. S1) used in this study for determining and quantifying threat magnitude, with some modifications described in the following paragraphs. This way, already existing Red List assessments (e.g., Norwegian Artsdatabanken (Artsdatabanken (2021))) could also be used.

The scoring matrix (Fig. S1) has the factors **scope** and **severity**; outside of the table, each scoring's **certainty** was also assessed. We have eliminated timing as a factor by only including threats that are currently ongoing; future threats are outside the scope of this study. However, it is worth noting that the impacts of many of the included threats are expected to increase in the future due to climate change and intensified human activity in the region (Smith & Stephenson (2013); VanWormer *et al.* (2019); Programme (2010); Onarheim & Årthun (2017); Directorate (2024); Last (2023)).

For both the factors scope and severity, the level "Unknown" was included to account for lack of data. IUCN also lists "Unknown" as a level, but it is not included in their scoring table. The level "Unknown" also relates to the certainty of the scoring. To minimize uncertainty, assessments were made conservatively. For example, if we were certain of the severity being at least Moderate, but some sources indicate that it might be Serious, we scored Moderate, not Unknown or Serious. Below, the factors and their levels are described in more detail.

**Scope**

Scope describes the fraction of the population affected by the threat. In this assessment, the "whole" population was the population in the Barents Sea region.

- Whole (>90%)
- Majority (50–90%)
- Minority (<50%)
- Unknown

**Severity**

Describes how severe the threat is to the population. IUCN scores severity based on the rate of population decline the threat causes/is expected to cause, e.g., "Causing or likely to cause very rapid declines (>30% of the population over 10 years or three generations; whichever is the longer)". For taxa-threat pairs where this level of detailed information was available (in practice mainly when a Red List assessment has been made), the assessment of severity could be done using IUCN's criteria. Otherwise, the assessment was based on the effects the threat has on population mortality and reproduction rates. For clarity, the levels have thus been given new names which focus less on speed.

- Major
  Causes/Is likely to cause large increases in mortality rate and/or causes/is likely to cause large reductions in reproduction rate.
- Moderate
  Causes/Is likely to cause moderate increases in mortality rate and/or causes/is likely to cause moderate reductions in reproduction rate.
- Minor
  Causes/Is likely to cause small increases in mortality rate and/or causes/is likely to cause small reductions in reproduction rate. *This could for example be a threat that lowers individual fitness but that is mostly non-lethal.*
- Unknown

**Certainty**

How certain the threat scoring is. The criteria below are applicable under the assumption that all literature used is credible and up to date.

- Certain
  A threat scoring was considered 'certain' if it was based on literature specifically about the taxon in question or a closely related taxon. "Closely related" means either taxonomically related or very similar in ecology and behavior. Which taxa can be considered "close enough" varies from case to case, so here the scorer needs to use their judgement and knowledge of the topic.

- Uncertain
  A threat scoring was considered 'uncertain' if there was insufficient data to score both factors, in other words, if the level of any factor was 'Unknown, *or* if the scoring was based on literature about a distantly related taxon (see point made above).

In the IUCN-scoring, the threats are initially scored (1–3) on the threat's timing. We removed timing as a factor, which would mean that a threat could get the score 0. However, we want 0 to mean that the threat is not acting upon a species — therefore we rescale so the lowest score (for the threat which scope and severity are both Unknown will be 1. This makes the range of impact scores in this scoring system (1–7) lower than in IUCN's (2–9).

We translated this values to an increased probability that a species goes extinct independent of its species interactions (i.e. presence/absence of preys).

| **Severity Scope** | Whole (3) | Majority (2) | Minority (1) | Unknown (0) |
|---|---|---|---|---|
| Severe (3) | 7 | 6 | 5 | 4 |
| Serious(2) | 6 | 5 | 4 | 3 |
| Moderate (1) | 5 | 4 | 3 | 2 |
| Unknown (0) | 4 | 3 | 2 | 1 |

Figure S1: Scoring table for threat impact.

#### Trophic level for threatened and service providing species

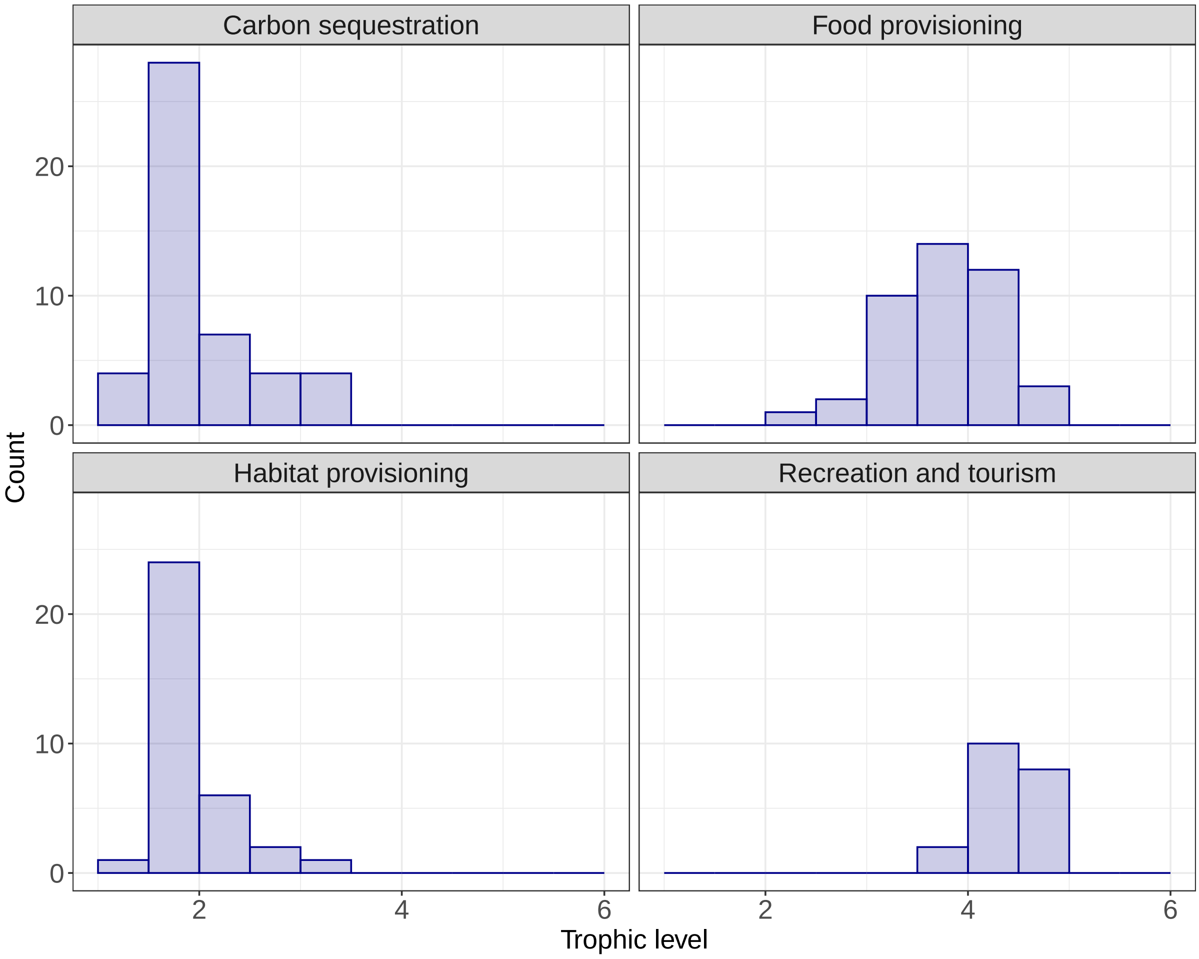


Figure S2: The histograms show the trophic levels (x-axis) for the species providing the respective service (panels). The y-axis shows the number of species for each respective trophic level. Note that the scale on the y-axis differ between panels. Trophic level is a continuous trait but has here been binned in steps of 0.25.

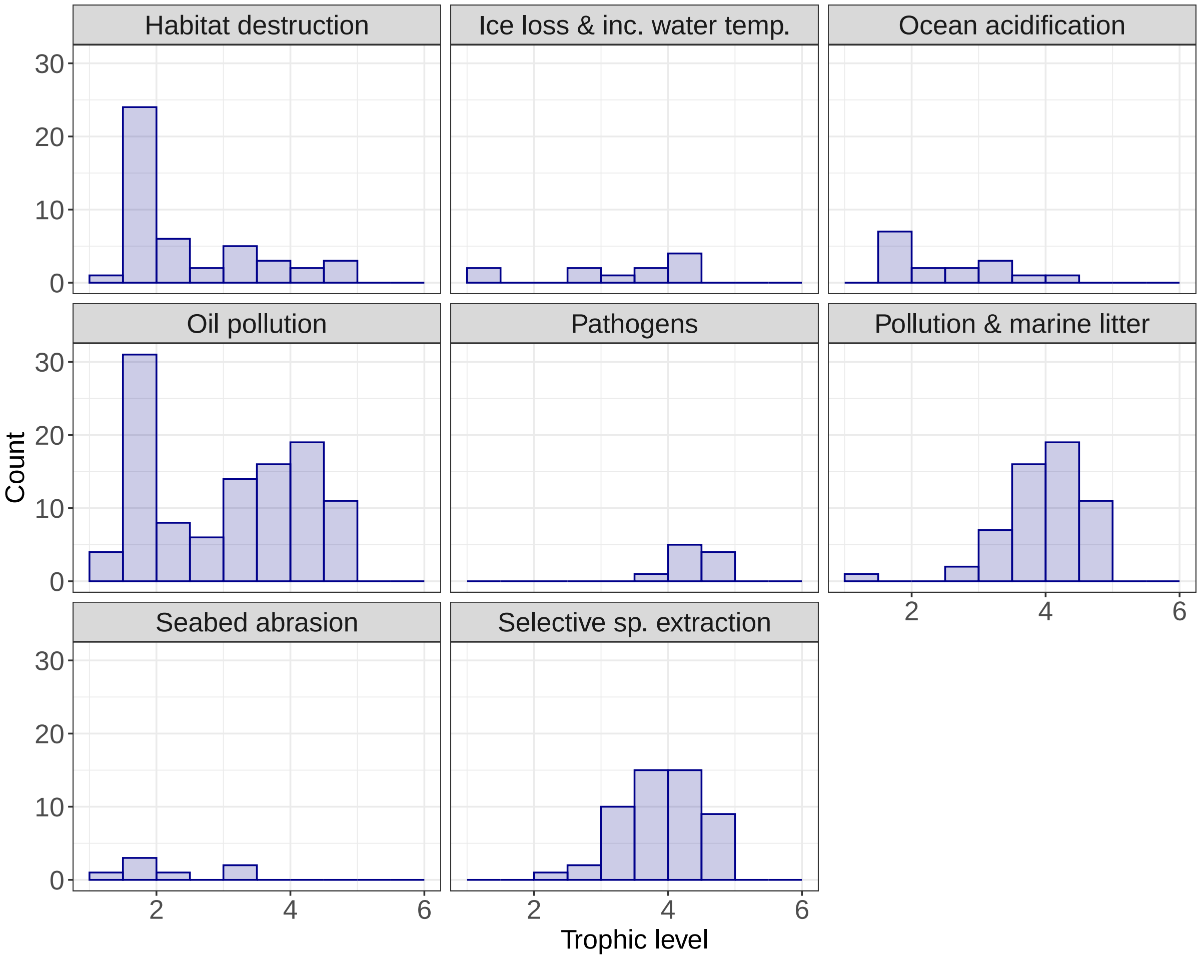


Figure S3: The histograms show the trophic levels (x-axis) for the species directly affected by respective threat (panels). The y-axis shows the number of species for each respective trophic level. Note that the scale on the y-axis differ between panels. Trophic level is a continuous trait but has here been binned in steps of 0.25.

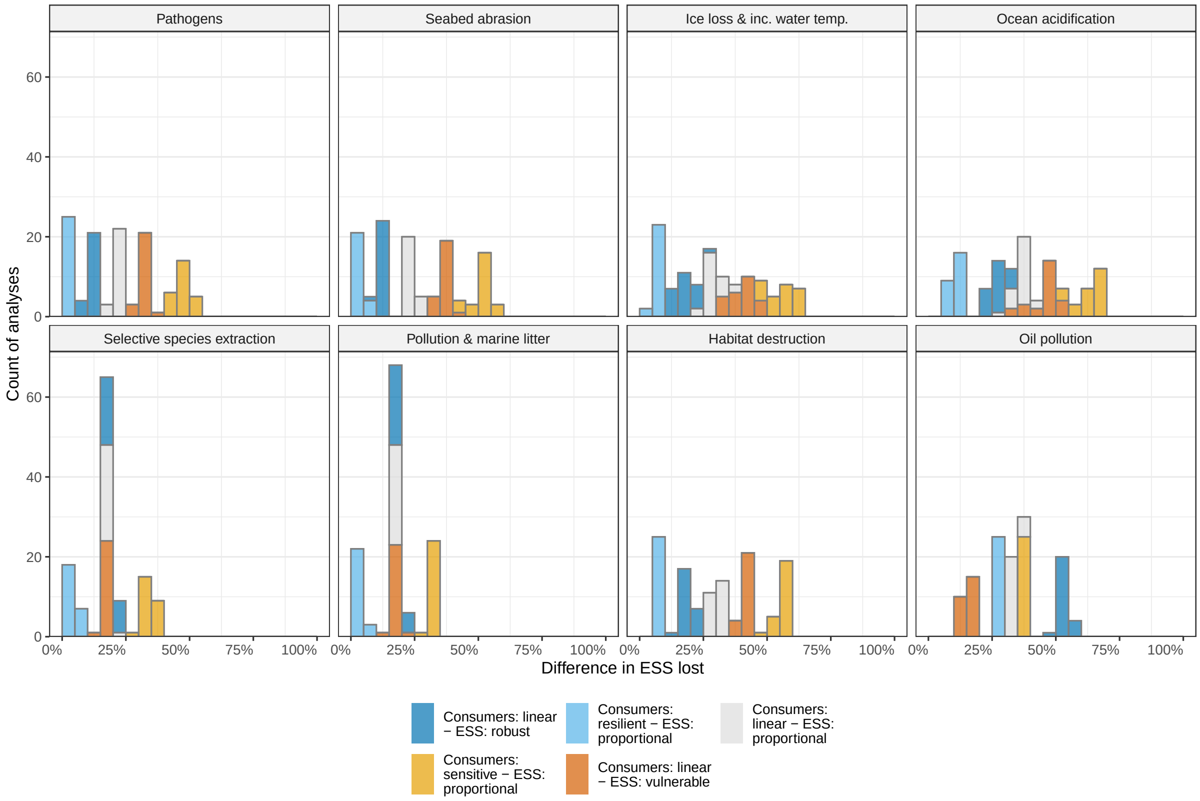


Figure S4: The number of analyses (y-axis) where indirect network effects caused increased loss of the ecosystem service Recreation and tourism after realization of different threats (panels). The x-axis shows the difference between ecosystem service retained when the species interactions are not taken into account versus when the interactions are taken into account, in percentages points. The histograms summarize calculations over the different subregions. Colors represent combinations of different consumer response and ecosystem service summary functions.

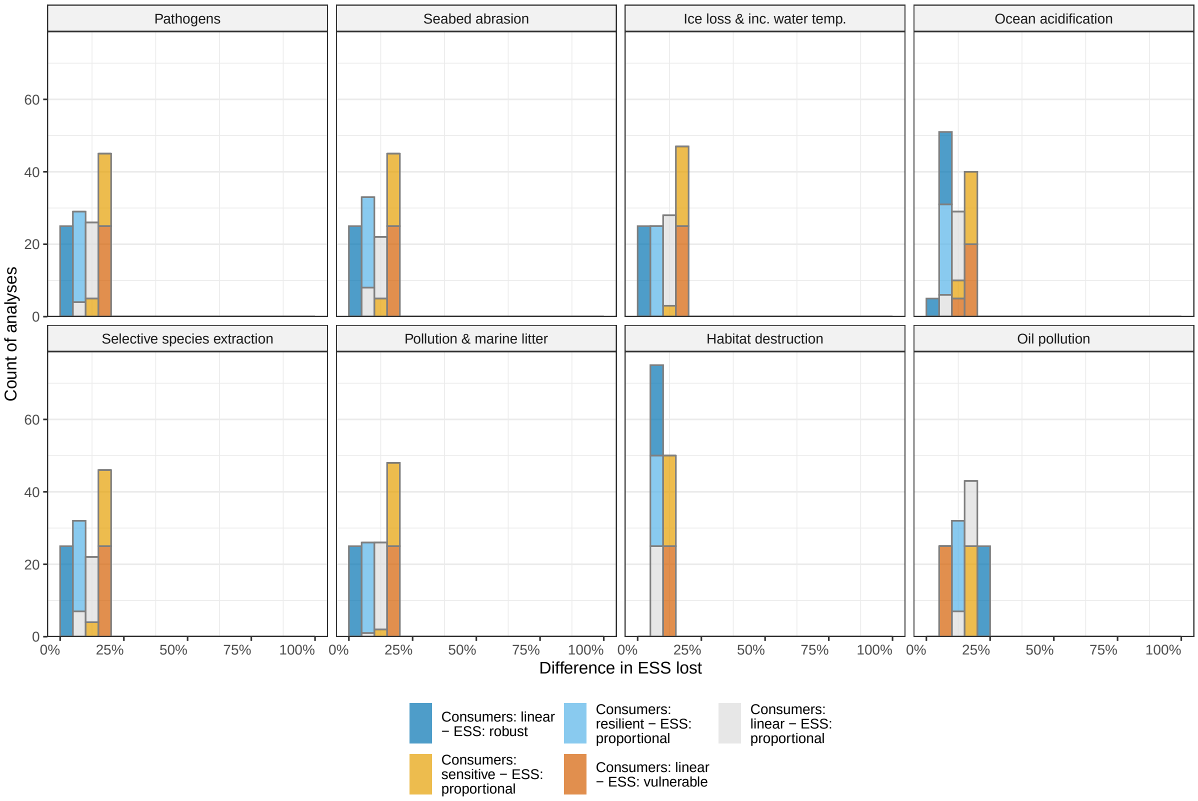


Figure S5: The number of analyses (y-axis) where indirect network effects caused increased loss of the ecosystem service Recreation and tourism after realization of different threats (panels). The x-axis shows the difference between ecosystem service retained when the species interactions are not taken into account versus when the interactions are taken into account, in percentages points. The histograms summarize calculations over the different subregions. Colors represent combinations of different consumer response and ecosystem service summary functions.

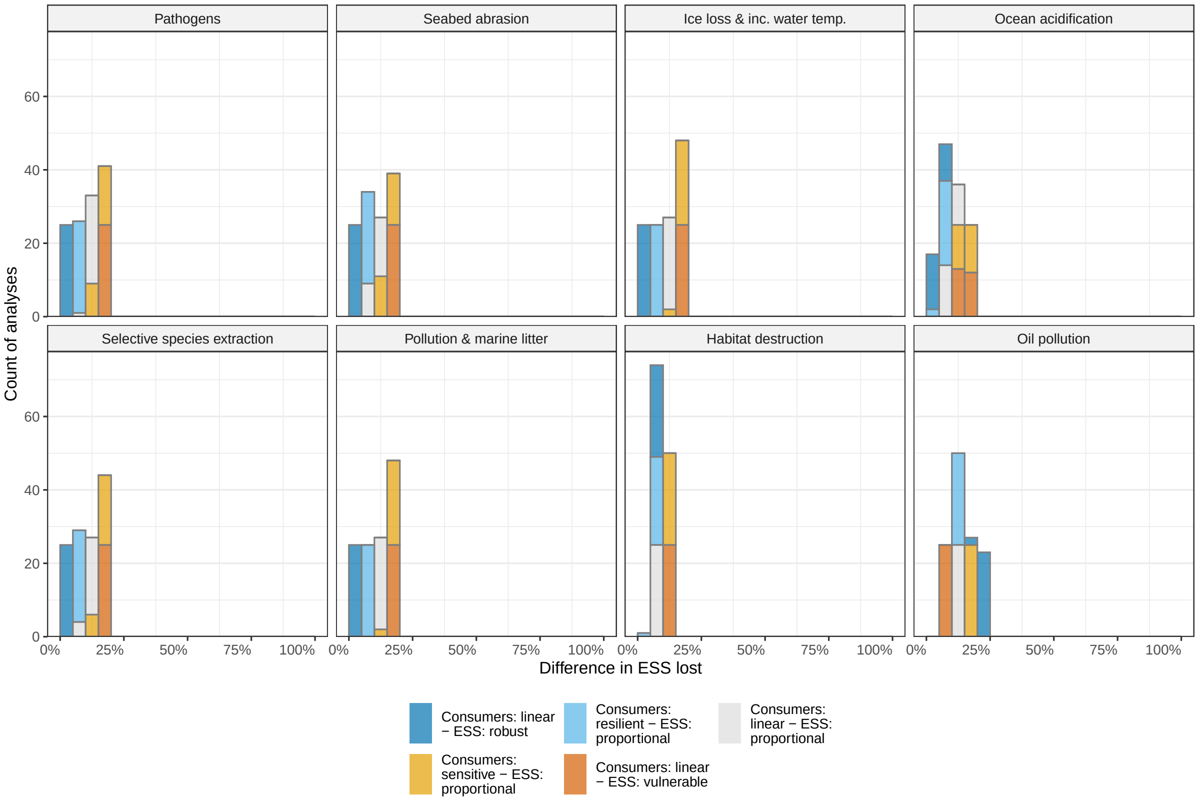


Figure S6: The number of analyses (y-axis) where indirect network effects caused increased loss of the ecosystem service Recreation and tourism after realization of different threats (panels). The x-axis shows the difference between ecosystem service retained when the species interactions are not taken into account versus when the interactions are taken into account, in percentages points. The histograms summarize calculations over the different subregions. Colors represent combinations of different consumer response and ecosystem service summary functions.

### References


AMAP (2013). AMAP Assessment 2013: Arctic Ocean Acidification. Tech. rep., Oslo, Norway.

AMAP (2018). AMAP Assessment 2018: Arctic Ocean Acidification. Tech. rep., Tromsø, Norway.

AMAP, A. (2010). Assessment 2007: oil and gas activities in the arctic-effects and potential effects. *Arctic Monitoring & Assessment Programme, Oslo, vols*, 1.

Anderson, O.R.J., Small, C.J., Croxall, J.P., Dunn, E.K., Sullivan, B.J., Yates, O. & Black, A. (2011). Global seabird bycatch in longline fisheries. *Endangered Species Research*, 14, 91–106.

Archer, L.C., Atkinson, S.N., Lunn, N.J., Penk, S.R. & Molnár, P.K. (2025). Energetic constraints drive the decline of a sentinel polar bear population. *Science*, 387, 516–521.

Artsdatabanken (2021). Rødlista 2021 - Artsdatabanken.

Atwood, T.B., Romanou, A., DeVries, T., Lerner, P.E., Mayorga, J.S., Bradley, D., Cabral, R.B., Schmidt, G.A. & Sala, E. (2024). Atmospheric CO2 emissions and ocean acidification from bottom-trawling. *Frontiers in Marine Science*, 10.

Bajzak, C., Hammill, M., Stenson, G. & Prinsenberg, S. (2011). Drifting away: implications of changes in ice conditions for a pack-ice-breeding phocid, the harp seal (pagophilus groenlandicus). *Canadian journal of zoology*, 89, 1050–1062.

Barratclough, A., Ferguson, S.H., Lydersen, C., Thomas, P.O. & Kovacs, K.M. (2023). A Review of Circumpolar Arctic Marine Mammal Health—A Call to Action in a Time of Rapid Environmental Change. *Pathogens*, 12, 937. Number: 7 Publisher: Multidisciplinary Digital Publishing Institute.

Barry, T., Berteaux, D. & Bültmann, H. (eds.) (2013). *Arctic Biodiversity Assessment: status and trends in Arcitc biodiversity.* The Conservation of Arctic Flora and Fauna, Akureyri, Iceland.

Bergmann, M., Collard, F., Fabres, J., Gabrielsen, G.W., Provencher, J.F., Rochman, C.M., van Sebille, E. & Tekman, M.B. (2022). Plastic pollution in the Arctic. *Nature Reviews Earth & Environment*, 3, 323–337. Number: 5 Publisher: Nature Publishing Group.

Bogstad, B., Prozorkevich, D., Gjøsæater, H., Russkikh, A., Dolgov, A., Prokopchuk, I., Dalpadado, P., Gordeeva, A., Rey, A., Fall, J. & Lindal Jørgensen, L. (2021). Causes of capelin stock fluctuations.

Boitsov, S., Jensen, H.K.B. & Klungsøyr, J. (2009). Natural background and anthropogenic inputs of polycyclic aromatic hydrocarbons (pah) in sediments of south-western barents sea. *Marine Environmental Research*, 68, 236–245.

Buhl-Mortensen, L., Ellingsen, K.E., Buhl-Mortensen, P., Skaar, K.L. & Gonzalez-Mirelis, G. (2016). Trawling disturbance on megabenthos and sediment in the Barents Sea: chronic effects on density, diversity, and composition. *ICES Journal of Marine Science*, 73, i98–i114.

Buhl-Mortensen, P. (2017). Coral reefs in the Southern Barents Sea: habitat description and the effects of bottom fishing. *Marine Biology Research*, 13, 1027–1040. Publisher: Taylor & Francis _eprint: https://doi.org/10.1080/17451000.2017.1331040.

Carmack, E. & Wassmann, P. (2006). Food webs and physical–biological coupling on pan-Arctic shelves: Unifying concepts and comprehensive perspectives. *Progress in Oceanography*, 71, 446–477.

Christiansen, J.S. & Reist, J.D. (2013). Fishes. In: *Arctic Biodiversity Assessment: status and trends in Arctic biodiversity.* The Conservation of Arctic Flora and Fauna, Akureyri, Iceland, pp. 192–245.

Collard, F. & Ask, A. (2021). Plastic ingestion by Arctic fauna: A review. *Science of The Total Environment*, 786, 147462.

Dalpadado, P., Arrigo, K.R., van Dijken, G.L., Skjoldal, H.R., Bagøien, E., Dolgov, A.V., Prokopchuk, I.P. & Sperfeld, E. (2020). Climate effects on temporal and spatial dynamics of phytoplankton and zooplankton in the Barents Sea. *Progress in Oceanography*, 185, 102320.

Dalpadado, P., Ingvaldsen, R.B., Stige, L.C., Bogstad, B., Knutsen, T., Ottersen, G. & Ellertsen, B. (2012). Climate effects on Barents Sea ecosystem dynamics. *ICES Journal of Marine Science*, 69, 1303–1316.

Dias, M.P., Martin, R., Pearmain, E.J., Burfield, I.J., Small, C., Phillips, R.A., Yates, O., Lascelles, B., Borboroglu, P.G. & Croxall, J.P. (2019). Threats to seabirds: A global assessment. *Biological Conservation*, 237, 525–537.

Directorate, N.O. (2024). APA 2023.

Duignan, P.J., Van Bressem, M.F., Baker, J.D., Barbieri, M., Colegrove, K.M., De Guise, S., De Swart, R.L., Di Guardo, G., Dobson, A., Duprex, W.P., Early, G., Fauquier, D., Goldstein, T., Goodman, S.J., Grenfell, B., Groch, K.R., Gulland, F., Hall, A., Jensen, B.A., Lamy, K., Matassa, K., Mazzariol, S., Morris, S.E., Nielsen, O., Rotstein, D., Rowles, T.K., Saliki, J.T., Siebert, U., Waltzek, T. & Wellehan, J.F.X. (2014). Phocine Distemper Virus: Current Knowledge and Future Directions. *Viruses*, 6, 5093–5134. Number: 12 Publisher: Multidisciplinary Digital Publishing Institute.

Edgar, G.J. (2025). Iucn red list criteria fail to recognise most threatened and extinct species. *Biological Conservation*, 301, 110880.

Fabry, V.J., Seibel, B.A., Feely, R.A. & Orr, J.C. (2008). Impacts of ocean acidification on marine fauna and ecosystem processes. *ICES Journal of Marine Science*, 65, 414–432.

Fossheim, M., Primicerio, R., Johannesen, E., Ingvaldsen, R.B., Aschan, M.M. & Dolgov, A.V. (2015). Recent warming leads to a rapid borealization of fish communities in the Arctic. *Nature Climate Change*, 5, 673–677. Number: 7 Publisher: Nature Publishing Group.

Fransner, F., Fröb, F., Tjiputra, J., Goris, N., Lauvset, S.K., Skjelvan, I., Jeansson, E., Omar, A., Chierici, M., Jones, E. *et al.* (2022). Acidification of the nordic seas. *Biogeosciences*, 19, 979–1012.

Freitas, F., Arndt, S., Hendry, K., Faust, J., Tessin, A. & März, C. (2022). Benthic organic matter transformation drives ph and carbonate chemistry in arctic marine sediments. *Global Biogeochemical Cycles*, 36, e2021GB007187.

Furness, R.W. (2003). Impacts of fisheries on seabird communities. *Scientia Marina*, 67, 33–45. Number: S2.

Gjøsæter, H., Huserbråten, M., Vikebø, F. & Eriksen, E. (2020). Key processes regulating the early life history of Barents Sea polar cod. *Polar Biology*, 43, 1015–1027.

Grøsvik, B.E., Prokhorova, T., Eriksen, E., Krivosheya, P., Horneland, P.A. & Prozorkevich, D. (2018). Assessment of Marine Litter in the Barents Sea, a Part of the Joint Norwegian–Russian Ecosystem Survey. *Frontiers in Marine Science*, 5.

Gundersen, H., Bryan, T., Chen, W., Moy, F.E., Sandman, A.N., Sundblad, G., Schneider, S., Andersen, J.H., Langaas, S. & Walday, M.G. (2017). *Ecosystem Services : In the Coastal Zone of the Nordic Countries*. Nordisk Ministerråd.

Halliday, W.D., Pine, M.K. & Insley, S.J. (2020). Underwater noise and Arctic marine mammals: review and policy recommendations. *Environmental Reviews*, 28, 438–448. Publisher: NRC Research Press.

Hesthagen, T., Wienerroither, R., Bjelland, O., Byrkjedal, I., Fiske, P., Lynghammar, A., Nedreaas, K. & Straube, N. (2021). Fisker: Vurdering av lodde Mallotus villosus for Norge.

Horvat, C., Jones, D.R., Iams, S., Schroeder, D., Flocco, D. & Feltham, D. (2017). The frequency and extent of sub-ice phytoplankton blooms in the Arctic Ocean. *Science Advances*, 3, e1601191. Publisher: American Association for the Advancement of Science.

Huserbråten, M.B.O., Eriksen, E., Gjøsæter, H. & Vikebø, F. (2019). Polar cod in jeopardy under the retreating Arctic sea ice. *Communications Biology*, 2, 1–8. Number: 1 Publisher: Nature Publishing Group.

ICES (2019). Barents Sea Ecoregion – Ecosystem overview. report, ICES Advice: Ecosystem Overviews.

ICES (2021a). Barents Sea Ecoregion – Ecosystem overview. report, ICES Advice: Ecosystem Overviews.

ICES (2021b). Barents Sea Ecoregion – Fisheries overview. report, ICES Advice: Fisheries Overviews.

ICES (2022). Barents Sea ecoregion – fisheries overview. report, ICES Advice: Fisheries Overviews.

IUCN (2023). Threats Classification Scheme (Version 3.3).

Ivanov, A.Y., Kucheiko, A.Y., Ivonin, D.V., Filimonova, N.A., Terleeva, N.V. & Evtushenko, N.V. (2022). Oil spills in the Barents Sea: The results of multiyear monitoring with synthetic aperture radar. *Marine Pollution Bulletin*, 179, 113677.

Johnston, D.W., Bowers, M.T., Friedlaender, A.S. & Lavigne, D.M. (2012). The effects of climate change on harp seals (pagophilus groenlandicus). *PLoS One*, 7, e29158.

de Jong, K., Forland, T.N., Amorim, M.C.P., Rieucau, G., Slabbekoorn, H. & Sivle, L.D. (2020). Predicting the effects of anthropogenic noise on fish reproduction. *Reviews in Fish Biology and Fisheries*, 30, 245–268.

Jørgensen, L.L., Planque, B., Thangstad, T.H. & Certain, G. (2016). Vulnerability of megabenthic species to trawling in the Barents Sea. *ICES Journal of Marine Science*, 73, i84–i97.

Koch, C.W., Brown, T.A., Amiraux, R., Ruiz-Gonzalez, C., MacCorquodale, M., Yunda-Guarin, G.A., Kohlbach, D., Loseto, L.L., Rosenberg, B., Hussey, N.E., Ferguson, S.H. & Yurkowski, D.J. (2023). Year-round utilization of sea ice-associated carbon in Arctic ecosystems. *Nature Communications*, 14, 1964. Number: 1 Publisher: Nature Publishing Group.

Krieger, K.J. & Wing, B.L. (2002). Megafauna associations with deepwater corals (Primnoa spp.) in the Gulf of Alaska. *Hydrobiologia*, 471, 83–90.

Kutti, T., Fosså, J.H. & Bergstad, O.A. (2015). Influence of structurally complex benthic habitats on fish distribution. *Marine Ecology Progress Series*, 520, 175–190.

Kühn, S., Bravo Rebolledo, E.L. & van Franeker, J.A. (2015). Deleterious Effects of Litter on Marine Life. In: *Marine Anthropogenic Litter* (eds. Bergmann, M., Gutow, L. & Klages, M.). Springer International Publishing, Cham, pp. 75–116.

Last, J. (2023). ‘Nature is being destroyed’: Russia’s arms buildup in Barents Sea creating toxic legacy. *The Guardian*.

Leu, E., Søreide, J.E., Hessen, D.O., Falk-Petersen, S. & Berge, J. (2011). Consequences of changing sea-ice cover for primary and secondary producers in the European Arctic shelf seas: Timing, quantity, and quality. *Progress in Oceanography*, 90, 18–32.

Lim, S.M., Payne, C.M., van Dijken, G.L. & Arrigo, K.R. (2022). Increases in Arctic sea ice algal habitat, 1985–2018. *Elementa: Science of the Anthropocene*, 10, 00008.

Lind, S., Ingvaldsen, R.B. & Furevik, T. (2018). Arctic warming hotspot in the northern Barents Sea linked to declining sea-ice import. *Nature Climate Change*, 8, 634–639. Number: 7 Publisher: Nature Publishing Group.

Madslien, K., Moldal, T., Gjerset, B., Gudmundsson, S., Follestad, A., Whittard, E., Tronerud, O.H., Dean, K.R., Åkerstedt, J., Jørgensen, H.J. *et al.* (2021). First detection of highly pathogenic avian influenza virus in norway. *BMC Veterinary Research*, 17, 218.

McBride, M.M., Hansen, J.R., Korneev, O. & Titov, O. (2016). Joint Norwegian - Russian environmental status 2013. Report on the Barents Sea Ecosystem. Part II - Complete report. Tech. Rep. 2.

Moore, S.E., Reeves, R.R., Southall, B.L., Ragen, T.J., Suydam, R.S. & Clark, C.W. (2012). A New Framework for Assessing the Effects of Anthropogenic Sound on Marine Mammals in a Rapidly Changing Arctic. *BioScience*, 62, 289–295.

Neimanis, A., Gavier-Widén, D., Leighton, F., Bollinger, T., Rocke, T. & Mörner, T. (2007). An outbreak of type c botulism in herring gulls (larus argentatus) in southeastern sweden. *Journal of wildlife diseases*, 43, 327–336.

Onarheim, I.H. & Årthun, M. (2017). Toward an ice-free Barents Sea. *Geophysical Research Letters*, 44, 8387–8395. _eprint: https://onlinelibrary.wiley.com/doi/pdf/10.1002/2017GL074304.

OSPAR (2008a). Coral gardens. Tech. rep.

OSPAR (2008b). Deep-sea sponge aggregations. Tech. rep.

OSPAR (2008c). Horse mussel beds. Tech. rep.

OSPAR (2009). Background Document for Lophelia pertusa reefs. Tech. rep.

OSPAR (2010). Background Document for Seamounts. Tech. rep.

OSPAR (2021a). Background document on kelp forest habitat. Tech. rep.

OSPAR (2021b). Case report for kelp forest habitat. Tech. rep.

OSPAR (2022a). Status Assessment 2022 - Coral Gardens.

OSPAR (2022b). Status Assessment 2022 - Lophelia pertusa reefs.

OSPAR (2024). List of Threatened and/or Declining Species & Habitats.

Programme, A.M.a.A. (2010). *AMAP Assessment 2007: Oil and gas activities in the Arctic - effects and potential effects. 2*. AMAP Arctic Monitoring and Assessment Programme, Oslo.

Rantanen, M., Karpechko, A.Y., Lipponen, A., Nordling, K., Hyvärinen, O., Ruosteenoja, K., Vihma, T. & Laaksonen, A. (2022). The Arctic has warmed nearly four times faster than the globe since 1979. *Communications Earth & Environment*, 3, 1–10. Number: 1 Publisher: Nature Publishing Group.

Rode, K.D., Regehr, E.V., Bromaghin, J.F., Wilson, R.R., St. Martin, M., Crawford, J.A. & Quakenbush, L.T. (2021). Seal body condition and atmospheric circulation patterns influence polar bear body condition, recruitment, and feeding ecology in the chukchi sea. *Global Change Biology*, 27, 2684–2701.

Rodolfo-Metalpa, R., Houlbrèque, F., Tambutté, , Boisson, F., Baggini, C., Patti, F.P., Jeffree, R., Fine, M., Foggo, A., Gattuso, J.P. & Hall-Spencer, J.M. (2011). Coral and mollusc resistance to ocean acidification adversely affected by warming. *Nature Climate Change*, 1, 308–312. Number: 6 Publisher: Nature Publishing Group.

Sala, E., Mayorga, J., Bradley, D., Cabral, R.B., Atwood, T.B., Auber, A., Cheung, W., Costello, C., Ferretti, F., Friedlander, A.M., Gaines, S.D., Garilao, C., Goodell, W., Halpern, B.S., Hinson, A., Kaschner, K., Kesner-Reyes, K., Leprieur, F., McGowan, J., Morgan, L.E., Mouillot, D., Palacios-Abrantes, J., Possingham, H.P., Rechberger, K.D., Worm, B. & Lubchenco, J. (2021). Protecting the global ocean for biodiversity, food and climate. *Nature*, 592, 397–402. Publisher: Nature Publishing Group.

Screen, J.A. & Simmonds, I. (2010). Increasing fall-winter energy loss from the Arctic Ocean and its role in Arctic temperature amplification. *Geophysical Research Letters*, 37. _eprint: https://onlinelibrary.wiley.com/doi/pdf/10.1029/2010GL044136.

Serreze, M.C. & Barry, R.G. (2011). Processes and impacts of Arctic amplification: A research synthesis. *Global and Planetary Change*, 77, 85–96.

Skagseth, Ø., Eldevik, T., Årthun, M., Asbjørnsen, H., Lien, V.S. & Smedsrud, L.H. (2020). Reduced efficiency of the barents sea cooling machine. *Nature Climate Change*, 10, 661–666.

Skogen, M.D., Olsen, A., Børsheim, K.Y., Sandø, A.B. & Skjelvan, I. (2014). Modelling ocean acidification in the Nordic and Barents Seas in present and future climate. *Journal of Marine Systems*, 131, 10–20.

Smith, L.C. & Stephenson, S.R. (2013). New Trans-Arctic shipping routes navigable by mid-century. *Proceedings of the National Academy of Sciences*, 110, E1191–E1195. Publisher: Proceedings of the National Academy of Sciences.

Tervo, O.M., Blackwell, S.B., Ditlevsen, S., Garde, E., Hansen, R.G., Samson, A.L., Conrad, A.S. & Heide-Jørgensen, M.P. (2023). Stuck in a corner: Anthropogenic noise threatens narwhals in their once pristine Arctic habitat. *Science Advances*, 9, eade0440. Publisher: American Association for the Advancement of Science.

VanWormer, E., Mazet, J.a.K., Hall, A., Gill, V.A., Boveng, P.L., London, J.M., Gelatt, T., Fadely, B.S., Lander, M.E., Sterling, J., Burkanov, V.N., Ream, R.R., Brock, P.M., Rea, L.D., Smith, B.R., Jeffers, A., Henstock, M., Rehberg, M.J., Burek-Huntington, K.A., Cosby, S.L., Hammond, J.A. & Goldstein, T. (2019). Viral emergence in marine mammals in the North Pacific may be linked to Arctic sea ice reduction. *Scientific Reports*, 9, 15569. Number: 1 Publisher: Nature Publishing Group.

Vongraven, D., Amstrup, S., McDonald, T., Mitchell, J. & Yoccoz, N. (2023). Relating polar bears killed, human presence, and ice conditions in svalbard 1987–2019. *Frontiers in Conservation Science*, 4, 1187527.

Wallhead, P.J., Bellerby, R.G.J., Silyakova, A., Slagstad, D. & Polukhin, A.A. (2017). Bottom Water Acidification and Warming on the Western Eurasian Arctic Shelves: Dynamical Downscaling Projections. *Journal of Geophysical Research: Oceans*, 122, 8126–8144. _eprint: https://onlinelibrary.wiley.com/doi/pdf/10.1002/2017JC013231.

Watelet, S., Skagseth, Ø., Lien, V.S., Sagen, H., Østensen, Ø., Ivshin, V. & Beckers, J.M. (2020). A volumetric census of the barents sea in a changing climate. *Earth System Science Data*, 12, 2447–2457.

Werner, S. & O'Brien, A.S. (2017). Marine litter. In: *Handbook on Marine Environment Protection: Science, Impacts and Sustainable Management*. Springer, pp. 447–461.

Yu, Y., Katsoyiannis, A., Bohlin-Nizzetto, P., Brorstrom-Lunden, E., Ma, J., Zhao, Y., Wu, Z., Tych, W., Mindham, D., Sverko, E. *et al.* (2019). Polycyclic aromatic hydrocarbons not declining in arctic air despite global emission reduction. *Environmental science & technology*, 53, 2375–2382.

Zhang, Z., Wu, P., Wang, X., Pang, Q., Wang, Y., Zhang, X., Kvale, K., Zeng, E.Y., Lei, L. & Zhang, Y. (2025). Ecological risk assessment of marine plastic pollution. *Nature Sustainability*, pp. 1–11.

Zhao, Y., Zhang, T., Lindenmayer, D. & Liu, J. (2025). Iucn red list underestimates national conservation needs of transboundary species. *Biological Conservation*, 312, 111517.

Žydelis, R., Bellebaum, J., Österblom, H., Vetemaa, M., Schirmeister, B., Stipniece, A., Dagys, M., van Eerden, M. & Garthe, S. (2009). Bycatch in gillnet fisheries – An overlooked threat to waterbird populations. *Biological Conservation*, 142, 1269–1281.
